# Egr-2 coordinates intestinal remodeling by promoting midline patterning and pharynx regeneration in planarians

**DOI:** 10.1101/2023.03.23.533990

**Authors:** Vasileios Morkotinis, David J. Forsthoefel

## Abstract

Regeneration requires remodeling of pre-existing structures and their integration with new cells and tissue. Identification of injury-induced regulators of these processes is central to regenerative biology. In planarian flatworms, the intestine remodels after amputation to re-establish normal morphology and integrate with the regenerating pharynx, the centrally located feeding organ. How these events are initiated and coordinated is currently unclear. In a candidate screen for injury-induced regulators of intestinal regeneration, we found that the transcription factor Early growth response protein 2 (Egr-2) was required for collective migration and fusion of intestinal branches at the anterior midline. Even though inhibition of *egr-2* by RNA interference increased dorsoventral (DV) muscle fibers at the midline between intestinal branches, bulk RNA sequencing of regenerating tissue revealed negligible dysregulation of DV-, body wall-, or intestinal-muscle-associated transcripts. Instead, the midline polarity cue *slit-1* and subsets of pharynx-associated transcripts were dysregulated. Both *egr-2* and *slit-1* RNAi delayed early pharynx regeneration, consistent with reduced *slit-1* expression in *egr-2(RNAi)* regenerates in the region of pharynx specification. Furthermore, inhibition of pharynx regeneration regulators *foxF-1* and *foxA* delayed integration with the intestine and disrupted formation of the intestine’s midline anterior branch. *Our findings suggest that the pharynx plays an inductive role in initiation of intestinal remodeling, and demonstrate how injury-induced reprogramming influences subsequent tissue assembly and integration during regeneration*.

## Introduction

Regeneration to repair or replace damaged body parts requires new tissue production. Substantial efforts in animal models are underway to understand the cellular origins of new tissue, and the coordination of proliferation and differentiation^1–3^. Less well understood, however, are the processes that occur simultaneously with the birth of new cells, including their assembly into tissues with three-dimensional structure, remodeling of pre-existing tissues, and integration of new and old tissues. Regeneration is initiated in part by transcriptional reprogramming in response to injury-induced signals^4, 5^. Changes in gene expression then coordinate de-differentiation, pluripotency, proliferation, patterning, and precisely timed differentiation of various tissue cell types^5–7^. Removing, ablating, or inhibiting differentiation of specific cell types^8–13^ can block subsequent proliferation or differentiation, but how cell-cell interactions induce later steps of tissue assembly has not been extensively investigated. Therefore, resolving how reprogramming links early wound signaling and new tissue production with remodeling and integration is a major goal.

The digestive system of planarian flatworms is a useful model for understanding regeneration at multiple levels of tissue organization. Planarians are capable of whole body regeneration, and restore all organs and tissues in about two weeks, even in tiny fragments^14, 15^. After surgical amputation, planarian tissue fragments regenerate the three primary branches of their intestine (one anterior, two posterior). This is achieved through differentiation of new intestinal cells by planarian adult stem cells called neoblasts, but also by extensive remodeling, collective migration, and merging of intestinal tissue that existed prior to injury^16–18^. In addition, small fragments lacking a centrally located feeding organ called the pharynx must completely regenerate this organ *de novo,* and the pharynx and intestine must re-integrate to establish a continuous lumen between the intestine and the pharynx’s esophagus^17, 19–22^. In tissue fragments from sublethally irradiated animals lacking neoblasts, the pharynx fails to regenerate, and intestinal remodeling is almost completely blocked^16, 17^. This raises the question whether production of new intestinal tissue, new pharynx tissue, or both are required to initiate or guide intestinal remodeling and integration with the new pharynx^17^.

How intestinal remodeling and pharynx regeneration are linked to early injury-responsive signaling is not well understood. Inhibition of axial polarity cues and regulators can alter re-establishment of intestinal branching^23–26^, cause regeneration of additional ectopic pharynges^27–30^, or block pharynx regeneration and intestinal remodeling altogether^31, 32^. However, because most polarity regulators affect multiple tissues, it is unclear whether pharynx and intestine respond directly to these cues, or influence each others’ regeneration. Similarly, although multiple injury-responsive transcription factors (TFs) are upregulated early during regeneration^33–40^, it is currently unknown whether these or other factors promote expression of TFs that play more specific roles in differentiation of intestinal (e.g., *gata4/5/6-1^4^*^1^) and pharynx (e.g., *foxA^2^*^2^) cells. Thus, whether early wound signals modulate reprogramming required for intestinal remodeling and pharynx regeneration remains a significant knowledge gap.

Early growth response (Egr) proteins are highly conserved Cys2-His2-fold-containing zinc-finger TFs that regulate embryonic development^42–45^, but also function^46^ in response to injury and during regeneration of the nervous system^47^, cardiovascular system^48^, muscle^49, 50^, and liver^44, 51^ in vertebrates. Egr TFs are widely conserved and are also upregulated by injury in organisms with greater regenerative capacity than mammals, including cnidarians^52^ and sea stars^53^. In acoels, Egr-1 functions as a pioneer TF, driving chromatin accessibility changes in response to amputation injury^54^. In planarians, at least six Egr proteins have been identified. Several are constitutively expressed, but most are upregulated by injury^34, 35, 37, 55, 56^. Egr-4 promotes formation of brain primordia and head regeneration^55^, while Egr-5 promotes differentiation of epidermal progeny cells^56^. Although functions for other Egr TFs have not been identified, the early tissue-specific expression and functions of *egr* family members in planarians and other animals suggested this family of proteins might have additional, unappreciated functions in regulating digestive organ regeneration.

Here, we identify a role for the Egr protein Egr-2 in intestine and pharynx regeneration in the planarian *S. mediterranea. egr-2* inhibition prevented fusion of intestinal branches at the anterior midline and disrupted integration with the pharynx in posterior fragments. Analysis of gene expression in regenerating tissue and genetic interactions with other regulators of muscle specification, midline polarity, and pharynx development revealed that *egr-2* regulated intestinal regeneration indirectly, by coordinating spatiotemporal re-expression of the midline polarity cue *slit-1* and promoting early pharynx regeneration. These observations demonstrate an inductive role for pharynx regeneration upstream of intestinal remodeling, an unexpected role for the midline repellent *slit-1* in pharynx regeneration, and a link between injury-induced reprogramming and planarian digestive organ regeneration.

## Results

### *egr-2* is required for intestinal remodeling

To identify *egr* family genes with potential roles in intestine regeneration, we first searched the planarian genome and several transcriptomes^57–59^ for *egr* homologs. Along with six known *egr* genes^34, 35, 55, 56, 60^, we identified three additional *egr* family members (*egr-6, egr-7,* and *egr-8*) that encoded the characteristic three C2H2-type zinc finger domains, and that were phylogenetically most similar to other Egr family members in planarians and other animals (Fig. S1A-C and Data S1). To determine whether *egr* transcripts were expressed in regions of intestine regeneration, we conducted colorimetric whole mount in situ hybridization ("WISH")^61, 62^. Consistent with previous reports^35, 37, 39, 55, 56^, we observed expression in wound-proximal tissue 1-24 hours after amputation (Fig. 1A and Fig. S2A, S2C-D), CNS (Fig. S2D), and epidermal progenitor cells/progeny (Fig. S2E). We also observed previously unappreciated expression in the pre-existing and regenerating nervous systems (Fig. S2G), and regenerating pharynx (Fig. S2A, S2D, S2F-H, and Fig. 1A).

**Figure 1.**
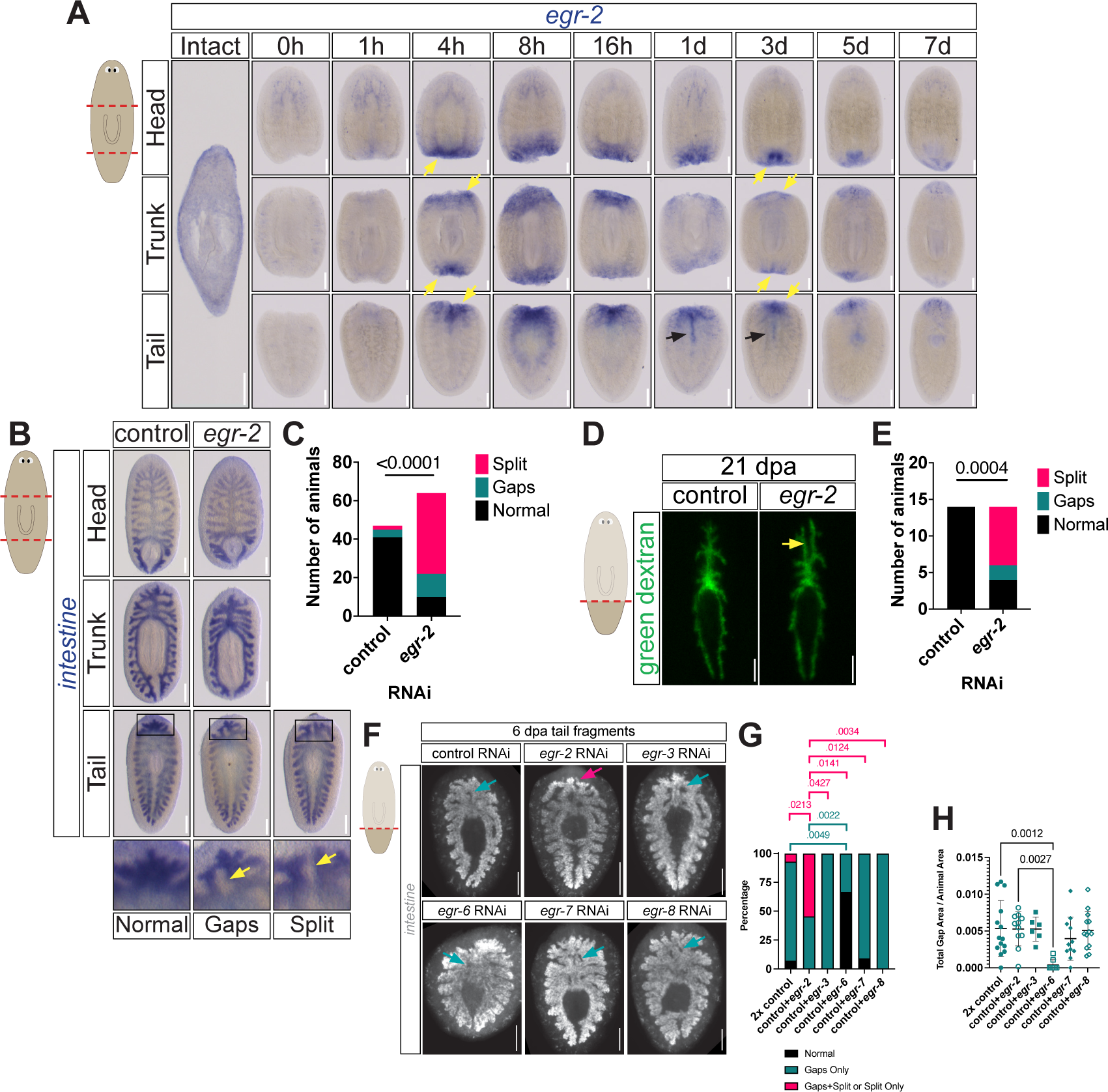
*egr-2* is required for intestinal remodeling. **(A)** Colorimetric ISH for *egr-2* mRNA over a 7 dpa time course. Schematic (left) shows amputation into head, trunk, and tail fragments. Yellow arrows, wound proximal regions of *egr-2* expression. Black arrows, midline *egr-2* expression in tail fragments. Dorsal view, anterior up. Scale bar, 200μm. **(B)** Colorimetric ISH with *apob-2* riboprobe labeling the intestine in tail regenerates at 6 dpa. Representative images of normal, gaps (arrow), and split gut (arrow) phenotypes. Dorsal view, anterior up. Scale bar, 200μm. **(C)** Quantification of *egr-2* RNAi phenotypes in (B). Chi-square test. **(D)** Intestine of live 21 dpa tail regenerates fed green fluorescent dextrans. Arrow: split anterior intestinal branches. Dorsal view, anterior up. Scale bar, 200μm. **(E)** Quantification of (D). Chi-square test. **(F)** Fluorescent ISH with intestine riboprobe pool on tail fragments (6 dpa) after single RNAi of *egr* genes. Magenta arrow, split gut. Teal arrows, gaps. Dorsal view, anterior up. Scale bar, 200μm. **(G)** Phenotype scoring in **(F)**. Fisher’s exact test, *P* values shown. **(H)** Normalized total gap area. Points indicate total gap area per animal divided by total animal area. Error bars, mean ±SD. Ordinary one-way ANOVA with Sídâk correction for multiple comparisons, *P* values shown.

*egr-2* was strongly upregulated in blastema-adjacent regions where both intestine remodeling and differentiation of new intestinal cells occur at later time points (3-7 dpa)^17^ (Fig. 1A). We used RNA interference (RNAi) to knock down *egr-2* (Fig. 1B and Fig. S3A) and evaluate its role in intestine regeneration. At 6 dpa, anterior fragments have normally regenerated two new posterior intestinal branches, and posterior fragments have regenerated a new primary anterior intestinal branch (Fig. 1B)^17^. In posterior fragments, *egr-2* RNAi disrupted the midline convergence of the two intestinal branches to re-establish the anterior primary intestinal branch (Fig. 1B-C). For example, in many fragments, the intestine’s connection to the regenerating pharynx was re-established, but branches failed to fuse at the midline more anteriorly, resulting in a forked or split appearance (Fig. 1B-C). In a higher proportion of *egr-2(RNAi)* animals vs. controls (12/64, ∼18.8% vs. 4/47, ∼8.5%), midline intestinal fusion occurred at the pharynx and also at anteriormost intestinal branches, but one or more gaps remained between these two regions, resulting in incomplete midline intestinal fusion (Fig. 1B-C). Intestinal regeneration in *egr-2(RNAi)* trunk fragments seemed to be unaffected, while in head fragments the distance between the tips of regenerating posterior primary gut branches was modestly increased, suggesting slightly slower growth (Fig. S3B-C). Together, these phenotypes suggested that egr-2 RNAi delayed intestine regeneration, and prevented attraction or collective migration of intestinal branches toward the midline.

To determine whether defects in intestine regeneration resolved at later timepoints, we examined intestinal morphology at 21 dpa by feeding high molecular weight fluorescent dextrans. These are selectively retained by intestinal phagocytes, enabling visualization of intestinal morphology in living regenerates^17, 63^. Anterior intestinal branch length was similar in control and *egr-2(RNAi)* animals (Fig. 1D), suggesting *egr-2* was not required for the intestinal growth or elongation. However, the split-gut phenotype was still present in over half of animals in posterior *egr-2(RNAi)* fragments, suggesting that the failure of intestinal branches to fuse at the midline was not a modest regeneration delay, but a persistent phenotype (Figure 1E).

Several other *egr* transcripts (*egr-3, egr-6, egr-7,* and *egr-8*) were also upregulated in regions of intestine regeneration (Fig. S2C and S2F-H). Using fluorescent in situ hybridization (“FISH”) to enable more sensitive detection and quantification of midline gaps in addition to anterior branch fusion defects, we found that knockdown of these other *egr* genes had only minor effects on intestinal regeneration. For example, we never observed the split-gut phenotype in *egr-3*, *egr-6*, *egr-7*, or *egr-8* single knockdowns (Fig. 1F-G). Furthermore, the number and relative total area of midline gaps were similar to control, except in *egr-6(RNAi)* fragments, in which the midline gap number and area were lower, suggesting *egr-6* might normally inhibit midline fusion (Fig. 1H). In double knockdowns with *egr-2*, *egr-3* RNAi modestly increased the proportion of animals with the split-gut phenotype (Fig. S3D-F). By contrast, the number of animals with gaps was modestly reduced in *egr-2;egr-6(RNAi*) fragments, further suggesting a possible inhibitory role for *egr-6* (Fig. S3D-F). Overall, these data suggested that other Egr TFs played redundant or more dispensable roles in anterior intestinal regeneration in tail fragments compared to *egr-2*.

*egr-2* RNAi did not affect animal growth (Fig. S4A-B), blastema size (Fig. S4C-D), pharynx regeneration (Fig. S4E-F), or brain regeneration (Fig. S4G-H), except in smaller *egr-2(RNAi)* tail fragments that had proportionally larger brains (Data S1). Eye regeneration was unaffected, and intereye distance and AP positioning was normal (Fig. S4I-K). Brain proportions were also normal, again with the exception of some small *egr-2(RNAi)* tail fragments (Fig. S4L-N and Data S1). Thus, *egr-2* RNAi did not seem to affect growth, regeneration, or scaling of other tissues. Altogether, these data indicated that *egr-2* was required more specifically for intestinal remodeling, especially in posterior fragments.

### Inhibition of midline dorsoventral muscle specification suppresses *egr-2* RNAi-induced intestinal defects

Although *egr-2* RNAi affected intestine remodeling, *egr-2* mRNA was strongly upregulated along the anterior midline in 1-3 dpa tail fragments (Fig. 1A), suggesting *egr-2* might function non-autonomously in adjacent midline tissues. Supporting this idea, numerous muscle fibers remained at the midline in 5 dpa *egr-2(RNAi)* fragments, but were absent in control fragments (Fig. 2A). At an earlier time point (3 dpa), we also found elevated numbers of neoblasts in this region (Fig. 2B), some of which expressed *mp-1,* a marker of both intestinal muscle (IM) and dorsoventral muscle (DVM)^64^ (Fig. 2C). By 4 and 8 dpa, the number of *mp-1-* positive cells was 2-3 times higher at the anterior midline in *egr-2(RNAi)* tail fragments (Fig. 2D-E). In uninjured animals, however, the number of midline muscle cells between posterior intestinal branches was comparable in control and *egr-2(RNAi)* animals (Fig. 2E). This suggested that Egr-2 might inhibit midline muscle birth or survival specifically during regeneration (Fig. 2E).

**Figure 2.**
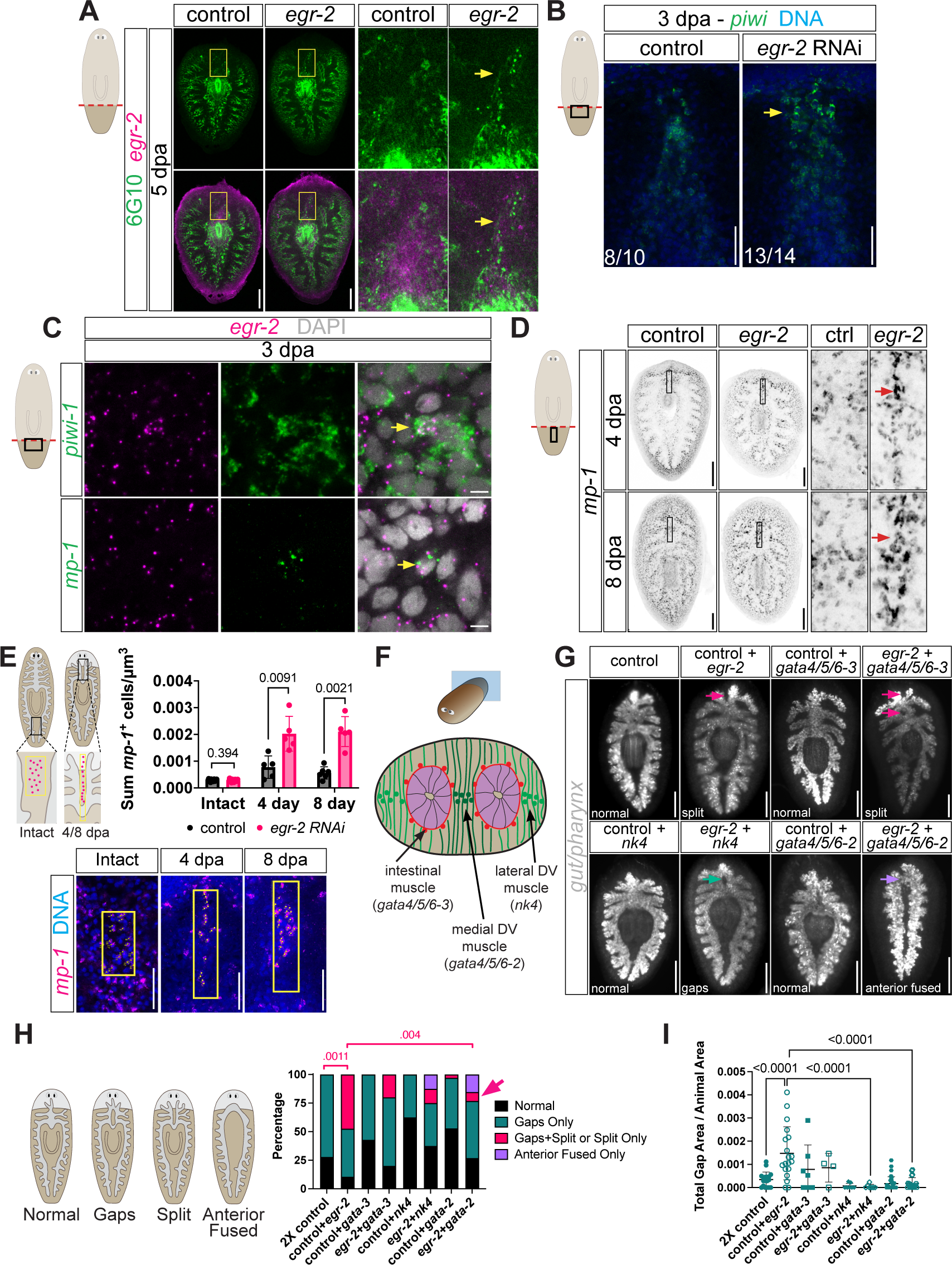
Inhibition of dorsoventral muscle specification suppresses *egr-2* RNAi phenotypes. **(A)** Fluorescent ISH for *egr-2* (magenta) and muscle fiber IF (anti-6G10, green) on 5 dpa tail fragments. Boxed regions at right. Arrows, ectopic midline muscle in *egr-2* RNAi fragments. Dorsal view, anterior up. Scale bar, 200 μm. **(B)** Fluorescent ISH with *piwi-1* riboprobe in 3 dpa tail fragments. Boxed region in schematic shown in images. Arrow, ectopic accumulation of neoblasts at anterior midline. Dorsal view, anterior up. Scale bar, 200 μm. **(C)** Co-expression (arrows) of *egr-2* (magenta) with *piwi-1* (green, upper panels) and *mp-1* (green, lower panels); fluorescent ISH in 3 dpa tail regenerates. Scale bar, 5 μm. **(D)** Fluorescent ISH for *mp-1^+^* muscle cells at 4 and 8 dpa. Inverted grayscale images. Boxed regions at right. Arrows, ectopic midline muscle cells in *egr-2* RNAi fragments. Dorsal view, anterior up. Scale bar, 200 μm. **(E)** Quantification of *mp-1^+^* cells between posterior gut branches in uninjured animals (left) and anterior gut branches (normalized to area scored) in 4 dpa and 8 dpa tail regenerates (right). Error bars, mean ±SD. Points represent data from individual animals. Lower panels, examples of regions scored. Welch’s *t* tests, *P* values shown. Scale bars, 50 μm. **(F)** Schematic of tail cross-section showing two posterior primary intestinal branches (purple), three non-body wall muscle cell types, and transcription factors required for their specification. **(G)** Fluorescent ISH with intestine riboprobe pool on 7 dpa tail fragments; RNAi of *egr-2* with muscle specifying transcription factors. Magenta arrows: split anterior intestinal branches; teal arrows: gaps. Dorsal view, anterior up. Scale bar, 200 μm. **(H)** Scoring of phenotypes in (G). Fisher’s exact test (*P* values shown for fragments with split or split plus gaps phenotypes). Fragment numbers with gaps only or anterior fusion were not significantly different. **(I)** Normalized total gap area for double knockdowns. Points indicate total gap area per animal divided by total animal area. Error bars, mean ±SD. Ordinary one-way ANOVA with Sídâk correction for multiple comparisons (*P* values shown).

To test whether midline muscle inhibited intestinal remodeling, we knocked down transcription factors required for specification of IM (*gata4/5/6-3*) and DVM (*nk4* and *gata4/5/6-2*)^64^ (Fig. 2F), individually and together with *egr-2*. Knockdown of these transcription factors individually had a negligible effect on anterior intestinal remodeling (Fig. 2G-H). Inhibition of IM specification by *gata4/5/6-3* RNAi modestly, but insignificantly suppressed the *egr-2* split-gut phenotype, and had no effect on the total area of midline intestinal gaps (Fig. 2G-I). By contrast, inhibition of medial DVM specification by knockdown of *gata4/5/6-2* reduced the number of animals with split anterior intestine and decreased the total midline gap area as compared to *egr-2* RNAi alone (Fig. 2G-I). Unexpectedly, inhibiting lateral DVM specification by *nk4* RNAi also modestly suppressed the split gut phenotype and significantly reduced midline gap area (Fig. 2G-I). Consistent with our results, inhibition of both IM and DVM specification caused midline fusion of posterior gut branches in uninjured animals, and reduced expression of the repulsive cue *slit-1* at the anterior midline in a previous study^64^. Thus, inappropriate midline muscle birth or survival after *egr-2* RNAi could delay intestine remodeling by secreting higher levels of midline repulsive cues, by structurally inhibiting collective migration of intestine branches towards the midline, or both.

Double RNAi with *egr-2* also caused a low penetrance phenotype in which the intestine only fused at its very anterior (4/26 *gata/4/5/6-2;egr-2(RNAi)* fragments and 1/9 *nk4;egr-2(RNAi)* fragments) (Fig. 2H). In addition, these fragments regenerated small or no pharynges (Fig. 2G). Although *gata4/5/6-2* RNAi causes uninjured animals to expel their pharynges^64^, a role in pharynx regeneration has not been reported for either *gata4/5/6-2* or *nk4.* Furthermore, we did not observe defects in pharynx regeneration in *egr-2(RNAi)* head or tail fragments at 6 dpa (Fig. S4E-F). Nonetheless, these rare phenotypes suggested a third possibility, that *egr-2* might promote an earlier step in pharynx regeneration, or a more upstream event that, when delayed, causes failed intestinal remodeling and inappropriate accumulation of midline DVM or IM cells.

### *egr-2* RNAi reduces expression of *sFRP-1, slit-1,* and injury-responsive, pharynx-associated transcripts

To distinguish between these possibilities, we first analyzed recently published scRNA-Seq data from regenerating tissue fragments^65^ to determine which cell types expressed *egr-2* (Fig. S5A-F). Consistent with a previous study^37^, *egr-2* was expressed in many cell types, including neoblasts, muscle, neurons, epidermis, and intestine (Fig. S5A-D). These included DVM, muscles expressing various polarity cues including *slit-1,* pharynx muscle (PM), and other pharynx cell types (Fig. S5E-F), suggesting that *egr-2* could function autonomously in both muscle and pharynx cell subpopulations.

Next, we performed bulk RNA sequencing (RNA-Seq) to assess global changes in gene expression in wound proximal regions of tail fragments at 0, 2, and 4 dpa (Figure 3A, Table S1), when transcripts involved in injury-responsive gene expression^35, 37^, re-establishment of axial polarity^66, 67^, neoblast proliferation^66–68^, and differentiation^68, 69^ are differentially expressed.

**Figure 3.**
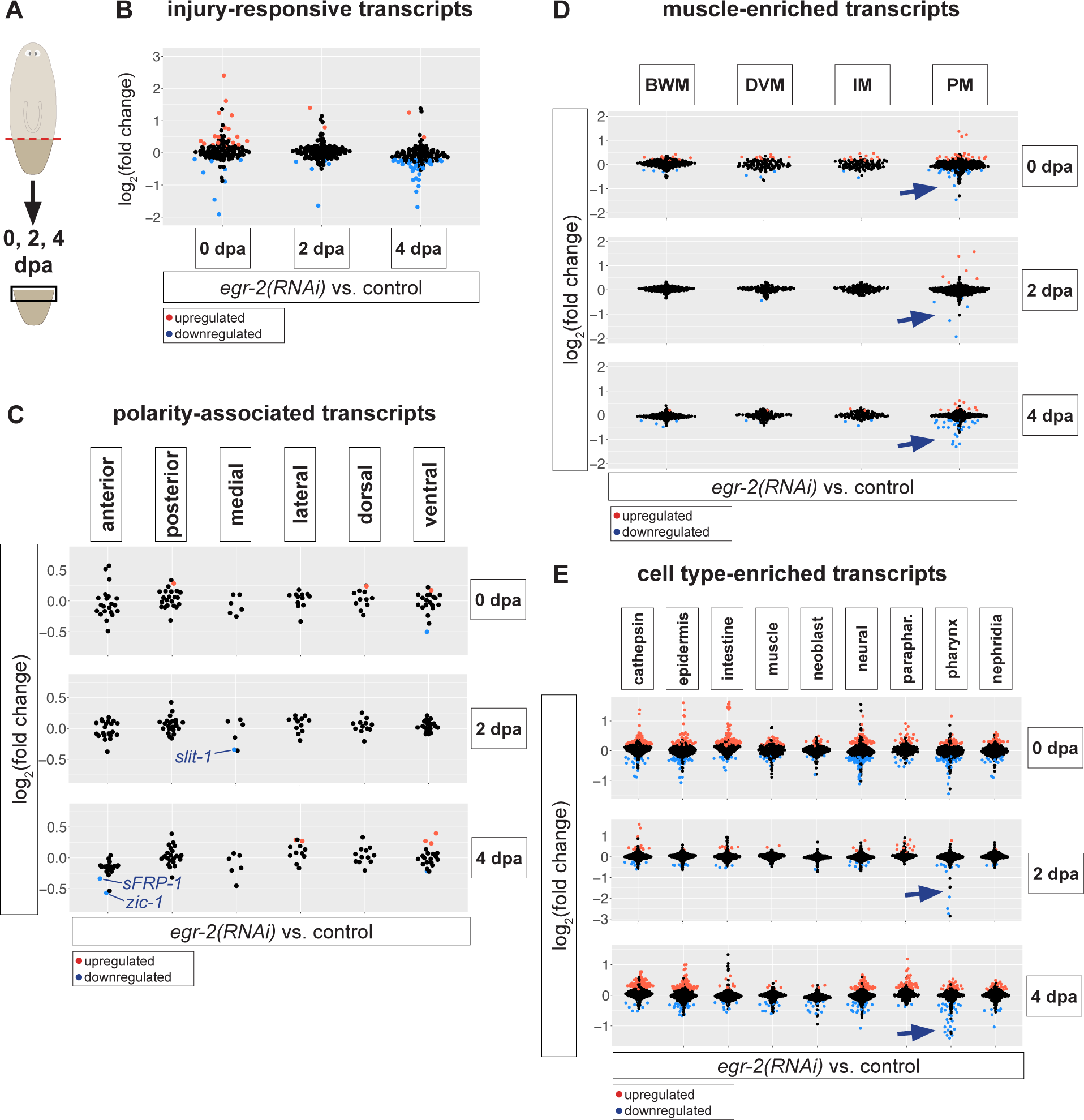
*egr-2* RNAi dysregulates polarity-and pharynx-related transcripts. **(A)** Schematic of tissue isolation strategy for RNA-Seq. **(B)** Differential expression of injury-responsive transcripts in *egr-2* RNAi vs. control RNAi wound-proximal tissue at the indicated time points. **(C)** Expression of polarity-associated transcripts in *egr-2* RNAi vs. control RNAi tissue. **(D)** Expression of muscle-enriched transcripts in *egr-2* RNAi vs. control RNAi tissue. Arrows, pharynx muscle-enriched transcripts downregulated by *egr-2* RNAi. BWM, body wall muscle. DVM, dorsoventral muscle. IM, intestinal muscle. PM, pharynx muscle. **(E)** Expression of cell type-enriched transcripts *egr-2* RNAi vs. control RNAi tissue. Arrows, pharynx-enriched transcripts downregulated by *egr-2* RNAi. In **(B-D)**, significantly up-or down-regulated transcripts (FDR-adjusted *P* value < 0.05) are shown in red or blue, respectively.

*egr-2* RNAi only modestly dysregulated the early transcriptional response to injury (Table S2A). Although over 100/128 injury-responsive transcripts^37^ were normally differentially expressed at 2 and 4 dpa in controls (vs. 0 dpa) (Fig. S6B), only seven transcripts were significantly dysregulated at 2 dpa, and 32 injury-responsive transcripts were downregulated at 4 dpa (Fig. 3B). Interestingly, when we cross-referenced these transcripts with a recent single cell analysis of planarian gene expression^70^, we found that 21/32 were enriched in muscle and/or pharynx cells (Fig. S6C), suggesting that *egr-2* might play a muscle-or pharynx-specific role in regulating injury-induced gene expression.

*egr-2* RNAi affected only a small subset of transcripts associated with anterioposterior (AP), dorsoventral (DV), and mediolateral (ML) identity, including many position control genes (PCGs) enriched in muscle and known to regulate axial polarity^37, 66, 67, 71^ (Fig. 3C, Fig. S6D, and Table S2B-D). Many of these transcripts were upregulated or downregulated in control tissue at 2 or 4 dpa (vs. 0 dpa) (Fig. S6D). However, only four transcripts were modestly dysregulated at 0 dpa, one transcript was downregulated at 2 dpa, and eight transcripts were dysregulated at 4 dpa (Fig. 3C). These included the repulsive midline cue *slit-1*^23^ and anterior-associated transcripts, *sFRP-1*^24, 25^ and *zic-1*^72^. With the exception of the HOX gene *post-2b*^73, 74^, which regulates tissue segmentation during asexual reproduction^74^, the other dysregulated polarity-related transcripts have no known functions. Thus, *egr-2* inhibition did not globally disrupt re-establishment of axial polarity, consistent with normal brain and eye regeneration in 6 dpa *egr-2(RNAi)* regenerates (Fig. S4G-H and S4I-N). However, the modest downregulation of *slit-1* and *sFRP-1* suggested possible roles for these cues during intestinal remodeling.

*egr-2* RNAi modestly dysregulated transcripts enriched in body wall (BWM), intestinal (IM), dorsoventral (DVM), and pharynx (PM) muscle subtypes^64^ at 0 dpa (Fig. 3D, Fig. S6E, and Table S3A-B). Surprisingly, although many muscle-associated transcripts were injury responsive (Fig. S6E), most transcripts dysregulated by 2 and 4 dpa were PM-enriched (Fig. 3D). At 2 dpa, 11/367 (∼3%) of PM transcripts were dysregulated, and at 4 dpa, 36/367 (∼10%, 10 upregulated transcripts and 26 downregulated transcripts) were significantly dysregulated (Fig. 3D). These data suggested that *egr-2* primarily regulated gene expression and/or differentiation of PM, but not other muscle subtypes. Furthermore, the fact that so few DVM-enriched transcripts were affected by *egr-2* inhibition (including dd_9910/*gata4/5/6-2*, dd_25344/*nk4*, and dd_6811/*mp-1*) suggested that ectopic midline DV muscle we observed in *egr-2(RNAi)* regenerates (Fig. 2D-E) was due to differences in location, rather than overall abundance.

*egr-2* RNAi also dysregulated transcripts enriched in other cell types^70^ (Fig. 3E, Fig. S6F, and Table S4A). Although many cell-type-enriched transcripts were injury-responsive (Fig. S6F) at both 2 and 4 dpa, *egr-2* RNAi dysregulated relatively few transcripts at 2 dpa (Fig. 3E). Only one neoblast-associated transcript was significantly downregulated at 2 dpa (Fig. 3E and Table S4C), consistent with the lack of blastema defects in *egr-2(RNAi)* regenerates (Fig. S4C-D). Strikingly, only pharynx-associated transcripts were downregulated more than 2-fold at 2 dpa (Fig. 3E). Similarly, pharynx-enriched transcripts were downregulated to a greater degree than for other cell types at 4 dpa (Fig. 3E and Table S4D). Moreover, when we limited our analysis to transcripts that were normally upregulated at 4 dpa, 22/73 (∼30.1%) of pharynx-enriched transcripts were downregulated by *egr-2* RNAi, and nearly half of these were downregulated over 2-fold (Fig. S6G and Table S4E). These included the pharynx/muscle marker *laminin* (dd_8356), the pharynx regeneration regulator *foxA* (dd_10718)^22^, markers of differentiating pharynx progeny (dd_554 and dd_561)^70^, and a marker of the pharynx’s connection to the body and intestine ("attachment site") (dd_372)^70^ (Table S4E). Thus, although *egr-2* did not affect pharynx size by 6 dpa (Fig. S4E-F), these results suggested that *egr-2* might regulate earlier stages of pharynx regeneration.

In summary, *egr-2* knockdown had limited effects on injury responsive gene expression and polarity re-establishment, and only moderate effects on gene expression by most cell types, including the intestine. However, downregulation of *sFRP-1, slit-1,* and pharynx-enriched transcripts suggested that *egr-2* might be required to re-establish expression of specific positional cues and/or promote early stages of pharynx development, either of which could subsequently be required for intestinal remodeling.

### sFRP-1 and Slit-1 play unexpected roles in anterior intestine regeneration

*sFRP-1, slit-1,* and *zic-1* were downregulated by *egr-2* RNAi (Fig. 3C). Because *egr-2* RNAi did not cause *zic-1-*associated phenotypes like delays in eye or brain regeneration (Fig. S4G-N) or dysregulation of anterior polarity cues aside from *sFRP-1*^72^ (Fig. 3C and Fig. S7A-B), we focused on whether *sFRP-1* or *slit-1* influenced intestinal remodeling. We first examined *sFRP-1* and *slit-1* expression in regenerating tails using colorimetric ISH (Fig. 4A-D). In 2 dpa *egr-2(RNAi)* fragments, *sFRP-1-*expressing cells had not coalesced at the anterior midline to the same extent as in controls, and occupied a greater area (Fig. 4A-B). This phenotype resolved by 4 dpa, although some fragments (4/5) had reduced *sFRP-1* expression in the regenerating pharynx (Fig. 4A). Similarly, *slit-1-*expressing cells also had not coalesced at the anterior midline in 2 dpa *egr-2(RNAi)* fragments (Fig. 4C-D), although the overall area occupied by *slit-1-*positive cells in the anterior 33% of the animal was similar (Fig. 4D). We also noticed elevated *slit-1* expression in the region of the regenerating pharynx in 4 dpa fragments (Fig. 4C). To test whether *egr-2* was required for early *slit-1* expression in this region, we used FISH to analyze expression quantitatively at early time points when the pharynx first begins to regenerate (Fig. 4E-F and Fig. S7C). Intriguingly, there was a significant reduction in *slit-1* signal in 2 dpa *egr-2(RNAi)* fragments that had resolved by 3 dpa (Fig. 4E-F and Fig. S7C), suggesting *egr-2* regulates this early subdomain of *slit-1* expression.

**Figure 4.**
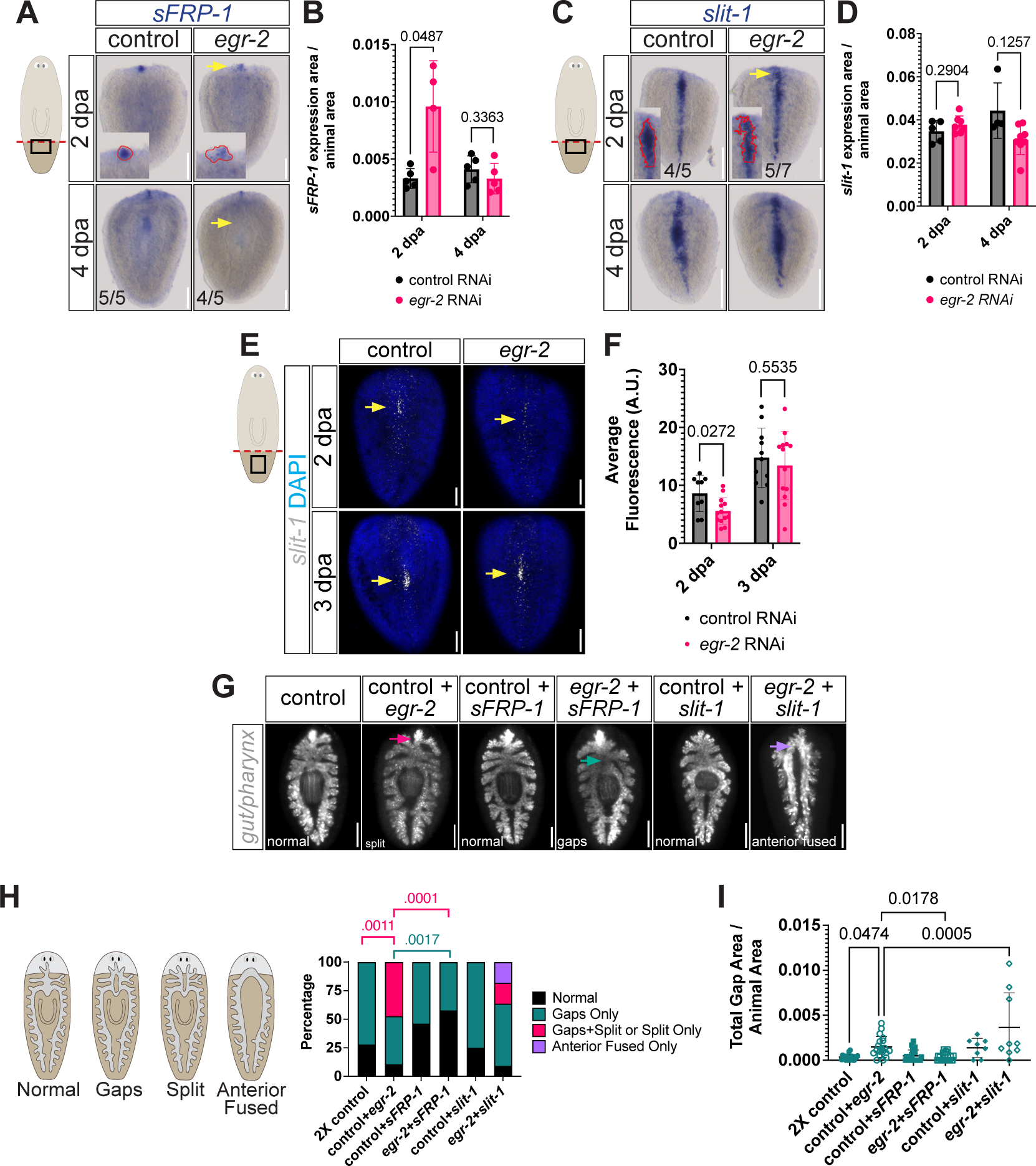
*slit-1* RNAi exacerbates intestine phenotypes in *egr-2* RNAi animals. **(A)** Colorimetric ISH for *sFRP-1* in 2 dpa and 4 dpa tail regenerates. Top arrow, example of delayed coalescence of anterior *sFRP-1+* cells (covering a larger region, see insets) in *egr-2* RNAi fragment. Bottom arrow, reduced *sFRP-1* expression anterior to the presumptive pharynx in *egr-2* RNAi fragments (4/5). Dorsal view, anterior up. Scale bar, 200 μm. **(B)** Quantification of normalized anterior *sFRP-1* expression area. Points represent data from individual animals. Error bars, mean ±SD. Welch’s *t* tests. **(C)** Colorimetric ISH for *slit-1* in 2 dpa and 4 dpa tail regenerates. Arrow, example of delayed coalescence of anterior *slit-1+* cells (5/7 *egr-2* RNAi fragments vs. only 1/5 control fragments). Insets, anterior *slit-1* expression area scored in **(D).** Dorsal view, anterior up. Scale bar, 100 μm. **(D)** Quantification of normalized anterior *slit-1* expression area. Points represent data from individual animals. Error bars, mean ±SD. Welch’s *t* tests. **(E)** Fluorescent ISH for *slit-1* (white) in 2 dpa and 3 dpa tail regenerates co-labeled with DAPI (blue). Ventral view, anterior up. Scale bar, 200 μm. Yellow arrows: *slit-1* expression quantified in (D) (see also Fig. S7C). **(F)** Quantification of *slit-1* fluorescence (arbitrary units, A.U.) in the ventral presumptive pharynx region, normalized to background fluorescence. Error bars, mean ±SD. Points represent values from individual animals. Welch’s *t* tests. **(G)** Fluorescent ISH with intestine/pharynx riboprobe pool in 7 dpa tail regenerates from single and double RNAi combinations. Magenta arrow, split anterior intestinal branches; teal arrow, gap; purple arrow, anterior intestinal fusion only. Dorsal view, anterior up. Scale bar, 200 μm. **(H)** Phenotype scoring from **(G)**. 2X control RNAi and control + *egr-2* RNAi data are identical to those in Figure 2G-I, because knockdowns were conducted as part of the same experiment. Fisher’s exact test, *P* values shown for conditions with significant differences in gaps only (teal) and split or split plus gap phenotypes. Fragment numbers with anterior fusion were not significantly different. **(I)** Normalized total gap area for double knockdowns. Ordinary one-way ANOVA with Šídâk correction for multiple comparisons, *P* values shown.

Since sFRP-1 is a putative inhibitor of Wnt signaling, its reduction in *egr-2(RNAi)* fragments might posteriorize tail fragments by decreasing antagonization of Wnt homologs secreted by posterior cells^16, 24, 25^, and cause the intestine to adopt a posterior-like, two-branched phenotype. Surprisingly, though, knockdown of *sFRP-1* suppressed the *egr-2* split gut phenotype, and reduced the total area of midline intestinal gaps (Fig. 4G-I). This suggested that sFRP-1 normally inhibits midline intestinal fusion, but also that sFRP-1’s downregulation by *egr-2* RNAi was not a primary cause of the split gut or gap phenotypes.

*slit-1* inhibition causes cyclopia, midline collapse of neural tissue during head regeneration, and midline fusion of posterior intestinal branches during tail regeneration, reflecting Slit-1’s role as a midline chemorepellent^23^. Thus *slit-1* inhibition might be expected to reduce midline repulsion and suppress *egr-2* RNAi’s disruption of midline intestinal fusion. Unexpectedly, however, *slit-1* RNAi caused a modest (but insignificant) increase in the total area of midline gaps in the regenerating anterior intestine compared to controls (Fig. 4I). Furthermore, although *slit-1* inhibition modestly reduced the number of animals with the split-gut phenotype in the *egr-2* background, the total area of midline gaps increased significantly when *slit-1* was knocked down together with *egr-2,* compared to *egr-2* RNAi alone (Fig. 4I). Additionally, in 2/11 *egr-2;slit-1(RNAi)* fragments, the intestine completely failed to fuse except at the very anterior of the fragment, accompanied by a complete lack of pharynx regeneration (Fig. 4G-I). *slit-1* was efficiently knocked down in these fragments, since 10/11 *slit-1(RNAi)* and 15/15 *egr-2;slit-1(RNAi)* fragments displayed cyclopia or narrower inter-eye distance (not shown). Together, these results suggested that *slit-1* either functioned oppositely in anterior tissue, as an intestinal chemoattractant, and/or that *slit-1* (together with *egr-2)* played an unappreciated role in pharynx regeneration, upstream of intestinal remodeling.

### *egr-2* and *slit-1* promote early pharynx regeneration

To test whether *egr-2* and *slit-1* might be required for earlier steps of pharynx regeneration, we first asked whether pharynx regeneration was affected by *slit-1* inhibition. As in our first analysis (Fig. S4E-F), pharynx size was unaffected in 6 dpa *egr-2(RNAi)* tail regenerates (Fig. 5A-B). However, in both *slit-1(RNAi)* and *egr-2;slit-1(RNAi)* 6 dpa regenerates, average pharynx size was reduced by ∼50% (Fig. 5A-B), suggesting that *slit-1* is required for efficient pharynx regeneration, and might function in parallel and/or downstream of *egr-2*.

**Figure 5.**
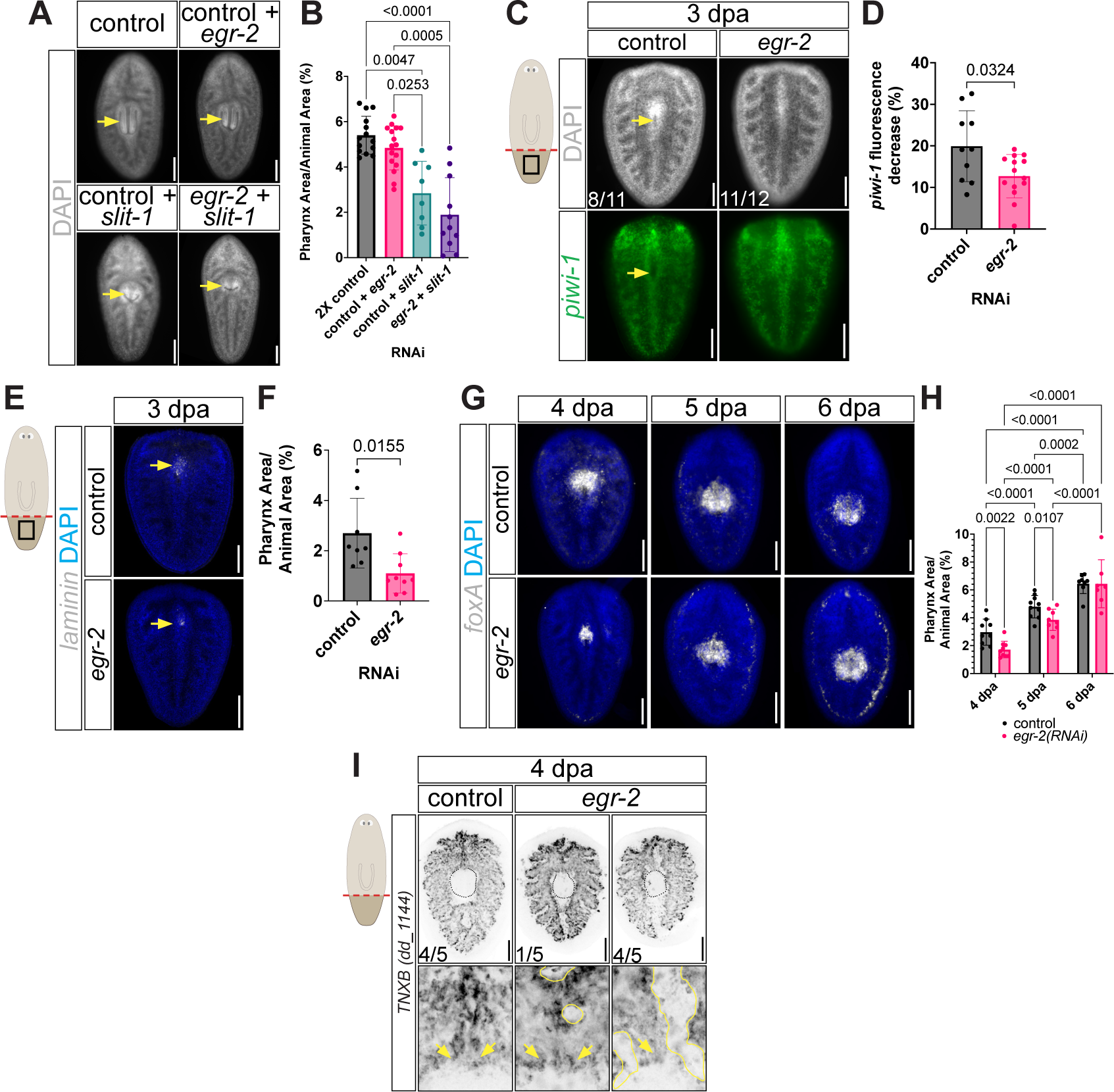
*egr-2* RNAi delays pharynx regeneration. **(A)** Pharynges (yellow arrows) in 7 dpa tail regenerates labeled with DAPI. Dorsal view, anterior up. Scale bar, 200 μm. **(B)** Normalized pharynx size in animals from **(A).** Error bars, mean ±SD. Points represent values from individual animals. Browne-Forsythe and Welch ANOVA with Dunnett’s T3 multiple comparisons test (significant *P* values shown). **(C)** Fluorescent ISH for *piwi-1* (green) in 3 dpa tail regenerates co-labeled with DAPI (gray). Yellow arrow: area of reduced DAPI and *piwi-1* expression indicative of pharyngeal cavity formation. Animal numbers with obvious pharyngeal cavity (control) or not (*egr-2*) in central DAPI-positive region are indicated. Dorsal view, anterior up. Scale bar, 200 μm. **(D)** Quantification of relative decrease in *piwi-1* fluorescence at the central midline/presumptive pharyngeal cavity in **(E)** (see also Figure S7D). Error bars, mean ±SD. Points represent values from individual animals. Welch’s *t* test, *P* value shown. **(E)** Fluorescent ISH for *laminin* (white) in 3 dpa tail regenerates co-labeled with DAPI (blue). Dorsal view, anterior up. Scale bar, 200 μm. Arrow, area of *laminin* expression quantified in (H). **(F)** Normalized pharynx area in 3 day tail regenerates. Points represent values from individual animals. Welch’s *t* test, *P* value shown. **(G)** Fluorescent ISH for *foxA* over a time course in tail regenerates. Dorsal view, anterior up. Scale bar, 200 μm. *foxA* expression area quantified in **(H)**. **(H)** Quantification of normalized pharynx size in **(G)**. Error bars, mean ±SD. Points represent values from individual animals. Two-way ANOVA with two-stage step-up method of Benjamini, Krieger, and Yekutieli correction for multiple comparisons (significant q values shown for comparisons between same times and conditions). **(I)** Fluorescent ISH for *TNXB* (dd_1144) in 4 dpa tail regenerates. Dotted lines (top images): pharynx perimeter. Yellow lines (bottom images): intestine gaps adjacent to pharynx (observed in 1/4 control fragments and 4/5 *egr-2(RNAi)* fragments). Arrows: intestine connection to esophagus/pharynx. Bottom images, magnified view of pharynx-intestine attachment site. Dorsal view, anterior up. Scale bar, 200 μm.

In uninjured planarians, the pharynx does not contain any *piwi-1^+^* neoblasts^75^ and thus, stem cells are gradually displaced from the central midline region as the pharyngeal cavity forms and pharynx regenerates. In 3 dpa regenerates, the formation of a DAPI-negative cavity in this region (8/11 control fragments) was reduced or absent in most *egr-2(RNAi)* fragments (11/12) (Fig. 5C). Quantitatively, the reduction of *piwi-1* signal in this region relative to immediately lateral regions was less pronounced compared to controls, indicating an early delay in the displacement of neoblasts and formation of the pharyngeal cavity (Fig. 5C-D and S7D).

At earlier time points, we found that *egr-2* RNAi also delayed pharynx regeneration. For example, the region of tissue expressing the pharynx muscle marker *laminin*^30^ was significantly smaller in 3 dpa *egr-2(RNAi)* fragments (Fig. 5E-F). Similarly, the area of regenerating pharynx expressing *foxA,* the earliest known regulator of pharynx regeneration^22^, was significantly smaller at 4 and 5 dpa, but not in 6 dpa fragments (Fig. 5G-H). Taken together, these observations suggested that *egr-2* was required for early pharynx regeneration, but that pharynx size is restored by later timepoints.

We predicted that an early delay in pharynx regeneration would also compromise early integration of intestinal branches with the newly formed pharynx. In 4 dpa control fragments, left and right portions of anterior intestinal branches had migrated to the midline just anterior to the regenerating pharynx (Fig. 5I), and midline gaps between intestinal branches were rare and generally less than 10 μm in width. In *egr-2(RNAi)* fragments, intestine was usually present immediately anterior to the pharynx, suggesting intestine-pharynx integration had partially occurred (Fig. 5I). However, in most fragments there were large adjacent gaps lateral to this primary branch (4/5 *egr-2(RNAi)* animals vs. 1/4 control animals) (Figure 5I). This suggested that an early delay in pharynx regeneration caused by *egr-2* RNAi could delay migration of intestinal tissue toward the midline, leading to persistent gaps and failed intestinal remodeling at later time points.

### Pharynx regeneration is required for efficient remodeling of the primary anterior intestine and triclad gut morphology

To further test the idea that pharynx regeneration was required for regeneration of the primary anterior intestinal branch, we performed additional amputation experiments. After midline amputation, sagittal *egr-2(RNAi)* fragments regenerated new posterior secondary branches adjacent to the pharynx, suggesting *egr-2* was dispensable for remodeling and differentiation in this region (Fig. S7E). After transverse amputation through the pharynx (Fig. S7F), anterior *egr-2(RNAi)* fragments regenerated posterior branches normally (not shown). Most posterior fragments, however, had obvious gaps just lateral to the pharynx attachment site (Fig. S7F-G), although the anterior branch was normal (not shown). Lastly, in thin transverse fragments from the post-pharyngeal region, *egr-2* RNAi caused the split intestine phenotype (Fig. S7H-I), increased total intestinal gap area (Fig. S7J), and reduced the size of the regenerated pharynx (Fig. S7K). However, in thin transverse regenerates from anterior regions, intestine regeneration was not impaired by *egr-2* RNAi (not shown), and pharynx regeneration was normal (Fig. S7L). Together, these experiments suggested that *egr-2* is mainly required for anterior intestinal remodeling when re-establishment of the primary anterior branch, *de novo* pharynx regeneration, and pharynx-intestine re-integration are all required.

We also directly tested the influence of pharynx on intestinal remodeling by knocking down two known regulators of pharynx regeneration. *forkhead box F-1 (foxF-1)* regulates specification of non-body wall muscle (IM, DVM, and PM)^64^, while *foxA* is currently the earliest known regulator of pharynx regeneration^22^. Consistent with previous studies^22, 64^, knockdown of both transcription factors strongly inhibited pharynx regeneration by 7 dpa (Fig. 6A-B). Likely because inhibition of *foxF-1* or *foxA* individually had such a severe effect, knockdown together with *egr-2* did not significantly enhance the disruption of pharynx regeneration (Fig. 6B).

**Figure 6.**
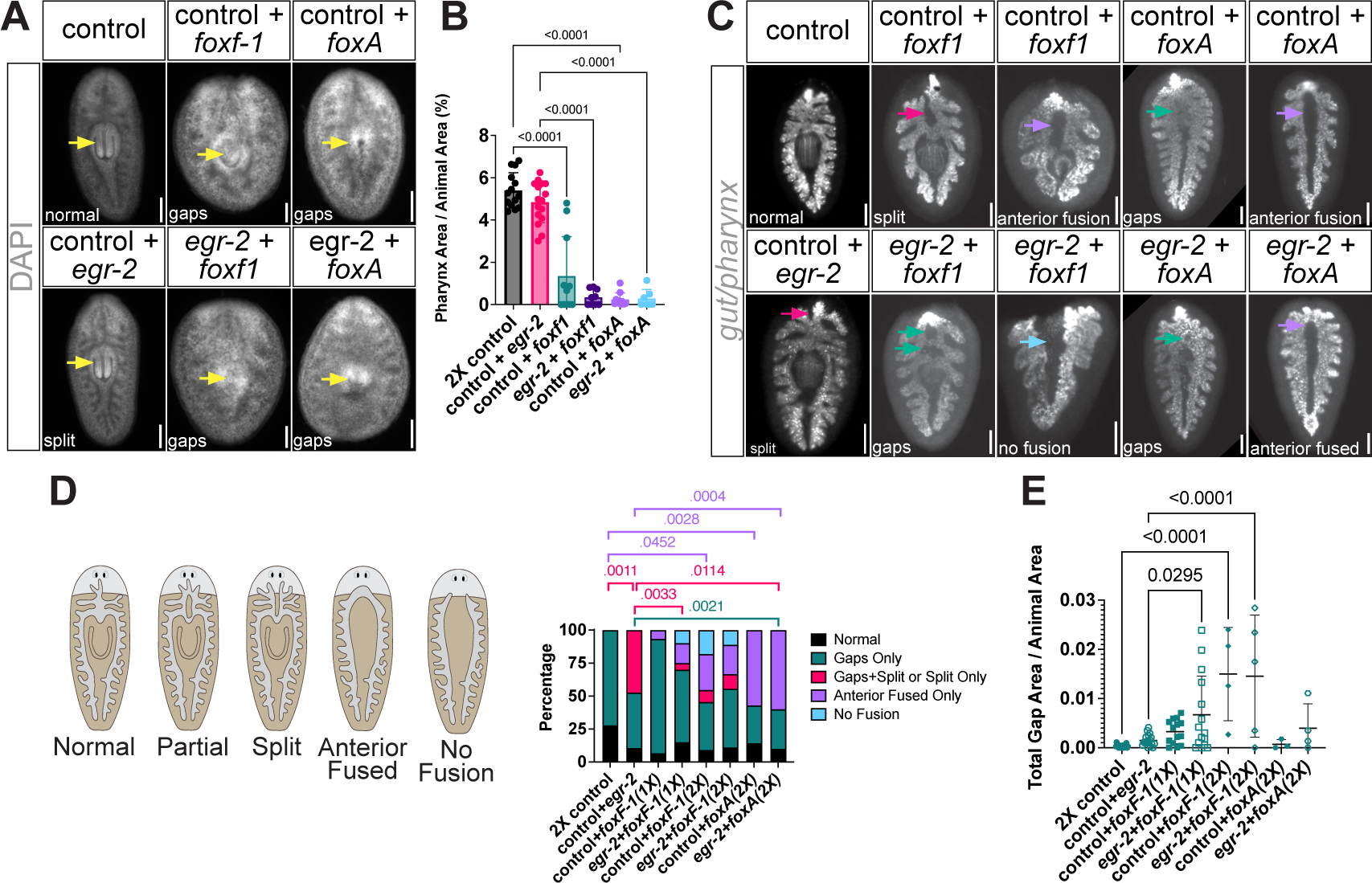
Pharynx regeneration is required for intestinal remodeling. **(A)** Reduced pharynx size in 7 dpa tail regenerates labeled with DAPI. Yellow arrows, pharynx. Dorsal view, anterior up. Scale bars, 200 μm. **(B)** Quantification of normalized pharynx size in animals from **(A)**. Dots represent values from individual animals. Error bars, mean ±SD. Ordinary one-way ANOVA with Šídák’s multiple comparisons test, *P* values shown. **(C)** Fluorescent ISH for intestine/pharynx riboprobe pool in 7 dpa tail regenerates from single and double RNAi combinations. Magenta arrow, split anterior intestinal branches. Teal arrow, gap. Purple arrow, anterior intestinal fusion. Dorsal view, anterior up. Scale bars, 200 μm. **(D)** Phenotype scoring from **(C)**. Fisher’s exact test, *P* values shown. Fragment numbers with anterior fusion were not significantly different. **(E)** Normalized total gap area for double knockdowns. Ordinary one-way ANOVA with Sídâk correction for multiple comparisons, *P* values shown. Animals were fed six doses of control or *egr-2* dsRNA; double knockdowns were also fed *foxF-1* or *foxA* either only for the last feeding (“1X”) or the last two feedings (“2X”). 2X control RNAi and control + *egr-2* RNAi data are identical to those in Figure 2G-I, because knockdowns were conducted as part of the same experiment.

*foxF-1* and *foxA* knockdown also severely disrupted anterior intestinal remodeling. Feeding even a single *foxF-1* dsRNA dose in an *egr-2(RNAi)* background significantly increased the total area of midline intestinal gaps relative to *egr-2* RNAi alone (Fig. 6C and Fig. 6E). Additionally, 3/20 *egr-2;foxF-1(RNAi-1X)* fragments displayed the “anterior fusion only/no pharynx” phenotype we also observed in *egr-2;slit-1(RNAi)* fragments (Fig. 4G-H), and 2/20 *egr-2;foxF-1(RNAi-1X)* fragments displayed a complete failure of intestinal fusion and no detectable pharynx regeneration (Fig. 6C-D). Feeding two *foxF-1* doses alone caused anterior fusion only and no fusion phenotypes, and increased gap area both alone and in combination with *egr-2* RNAi (Fig. 6C-E). Similarly, *foxA* RNAi (two doses) also caused the intestine to fuse only at its very anterior along with no detectable pharynx regeneration in 4/7 *foxA(RNAi-2X)* fragments and 6/10 *egr-2;foxA(RNAi-2X)* fragments (Fig. 6C-D). In the 3/4 remaining *egr-2;foxA* double knockdown fragments, total midline gap area was also modestly (but insignificantly) greater compared to *egr-2* or *foxA* single RNAi (Fig. 6E). Together, these experiments directly demonstrate that inhibiting pharynx regeneration disrupts anterior intestinal remodeling, supporting a primary role for *egr-2* as an early regulator of pharynx regeneration, which is itself an upstream requirement for intestinal regeneration.

## Discussion

A central problem in regeneration is how injury-induced transcriptional reprogramming coordinates the sequence of events required for successful re-establishment of organ morphology and function. Here, we found that the wound-responsive TF Egr-2 is required specifically for upregulation of *slit-1* at the central midline and pharynx regeneration in posterior regeneration fragments. Timely initiation of both events is required for collective migration of intestinal branches and their integration with the regenerating pharynx. suggest inductive roles for both pharynx and *slit-1,* and that *egr-2* likely regulates both cell autonomous and non-autonomous events required for pharynx and intestinal regeneration. More broadly, Egr-2’s role illustrates the importance of understanding how early signaling is linked through reprogramming factors to specific cell and tissue interactions upstream of remodeling and integration.

Our results indicate that several discrete steps are required for regeneration of the anterior intestine in posterior fragments (Fig. 7A). First, *slit-1* expression is re-established in a central midline domain where pharynx regeneration is initiated. Next, the anteriormost regions of the formerly posterior primary intestinal branches collectively migrate towards the midline, which occurs simultaneously with early pharynx regeneration. Subsequently, intestinal branches integrate with pharyngeal tissue at the pharynx attachment site, and fuse with each other at multiple more anterior locations. Finally, intestinal gaps are gradually resolved as midline muscle and other tissue is displaced or resorbed, and remaining intestine fuses to re-establish the primary anterior branch.

**Figure 7.**
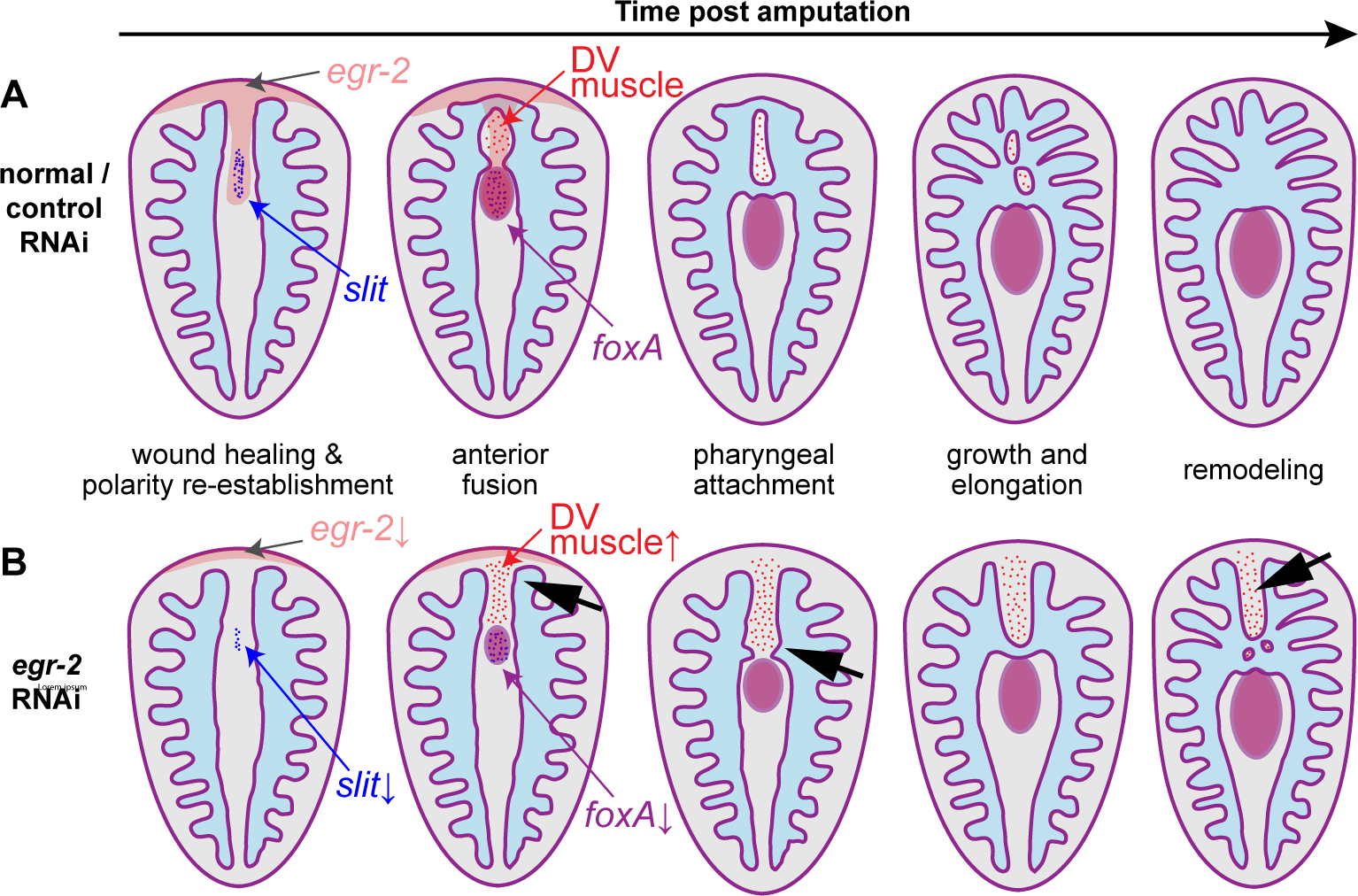
Model of anterior intestinal remodeling and Egr-2 function. **(A)** Normal timeline of *egr-2* expression (pink), *slit-1* expression (blue, midline patterning and pharynx induction), *foxA* expression (purple, pharynx progenitors and pharynx), and DV muscle differentiation/persistence (red) during remodeling of the intestine (light blue). **(B)** After *egr-2* RNAi, *slit-1* expression is reduced (blue), regeneration of *foxA+* pharynx tissue is delayed (purple), and DV muscle (red) accumulates. This delays anterior intestinal fusion, intestine attachment to the pharynx, and intestine remodeling to re-establish the primary anterior branch (black arrows).

Egr-2 could regulate pharynx regeneration and intestinal remodeling in multiple ways (Fig. 7B). In the most straightforward model, Egr-2 is required cell-autonomously for differentiation of central *slit-1* expressing muscle, as well as pharyngeal muscle and other pharynx cell types. *slit-1’s* expression at the central midline could promote mouth formation, invagination of the pharyngeal cavity, or migration of other muscles or progenitors required for early pharynx regeneration, although future experiments will be needed to distinguish *slit-1’s* precise roles. Interestingly, ectopic expression of the *Drosophila* Egr ortholog Stripe induces expression of Slit^76^. Inhibiting *egr-2* could thus delay pharynx regeneration both indirectly (by reducing *slit-1* expression) and directly (by delaying production of new pharynx cell types, which then delays medial migration of intestine branches). Delaying pharynx regeneration by either or both mechanisms would then delay medial migration of intestinal branches and integration with the pharynx. Additionally, this delay likely allows ectopic birth of new midline muscle, further inhibiting midline intestinal fusion.

Some of our observations challenge this simple model and suggest additional complexity. First, there was considerable variability in the penetrance and expressivity of intestinal phenotypes in *egr-2(RNAi)* fragments, and in gene interaction experiments. For example, we only observed the split gut phenotype in 30-50% of *egr-2(RNAi)* regenerates, and the number and total size of midline intestine gaps also varied. However, in previous studies^17, 77^, the number and location of anterior midline intestinal fusions (by 5 dpa) varied between individual fragments. In the present study, pharynx regeneration rate also varied across fragments. Thus, normal stochasticity in intestinal remodeling and pharynx regeneration could also explain the variability in the *egr-2* RNAi phenotype. Second, although intestinal fusion at the central midline was disrupted in 25-50% of *foxF-1(RNAi)* and *foxA(RNAi)* fragments, even in the complete absence of a new pharynx, the intestine usually fused at its anteriormost tips, suppressing the split gut phenotype. This could reflect *foxF-1’s* role in promoting *slit-1* expression in this anterior domain^64^; such a role could also be possible for *foxA,* although this has not yet been reported. Alternatively, intestinal remodeling might simply be able to occur independently of pharynx regeneration when midline fusion occurs more anteriorly, farther from the site of pharynx regeneration.

Egr-2 may also have additional roles. For example, over 100 transcripts normally enriched in *cathepsin+* cells were dysregulated in 2-4 dpa *egr-2(RNAi)* regenerates (Table S4). Because some *cathepsin+* cells are phagocytic^64^, *egr-2* inhibition could compromise their ability to clear dying midline cells or cellular debris in early regenerates, thereby inhibiting anterior intestinal fusion. As another example, Egr-2 might also control the influence of extracellular matrix or secreted cues that influence remodeling, since numerous proteases are dysregulated in *egr-2(RNAi)* regenerates (Table S1). Intriguingly, the *Drosophila* metalloprotease Tolkin cleaves Slit and converts it from a repulsive to a growth-promoting ligand^78^. A similar reduction in post-translational Slit-1 processing in *egr-2(RNAi)* regenerates could increase, rather than decrease, Slit-1-mediated repulsion, inhibiting midline fusion.

Current knowledge of wound-induced gene regulatory networks (GRNs) that promote successful regeneration is limited, in planarians as well as other organisms^5, 7^. In some animal models, broad ERK activation is induced by injury^33, 39, 79–81^, but how ERK and other signaling is ultimately transduced to coordinate cell-type-specific reprogramming is not well understood. Early *egr-2* upregulation in planarians is dependent on both ERK signaling^39^ and the MAP kinase activator *son-of-sevenless*^35^. Intriguingly, pharynx regeneration and intestine remodeling are also ERK-dependent^18^, as is expression of *foxA*^82^. We found *foxA* is downregulated by *egr-2* RNAi, suggesting an upstream link between early wound signaling and pharynx regeneration. More broadly, Egr proteins in numerous organisms have been implicated in regeneration of muscle and other tissues^44, 49, 53–55, 83, 84^, but their precise roles remain largely uncharacterized. Given that regeneration-specific enhancers drive tissue-specific gene expression^85, 86^, and that introduction of exogenous reprogramming factors promotes tissue repair^87–89^, further investigation of Egr TFs will be important to expand our understanding of how to manipulate injury-responsive GRNs.

Compared to recent progress in understanding the regulation of proliferation and differentiation during regeneration, how subsequent steps of tissue assembly and integration are orchestrated is less clear. During embryogenesis and regeneration, inductive cell-cell interactions promote subsequent development of adjacent tissues^90–96^. Our data suggest that the pharynx primordium plays an inductive role essential for intestine remodeling during planarian regeneration. An important next step will be to determine which nascent pharynx cell types promote remodeling, and whether this is achieved through secretion of pharynx-enriched signals^16^, exertion of adhesive or mechanical pulling forces between pharynx and intestine, or other mechanisms. In addition, our results do not formally exclude a reciprocal role for the intestine in regulating pharynx regeneration. For example, ectopic pharynges in *ptk7;wntP-2(RNAi)* and *ptk7;ndl-3(RNAi)* regenerates connect to intestinal tail branches with the pharynx-intestine attachment facing laterally, rather than anteriorly, suggesting the intestine influences pharynx positioning^29^.

Understanding how transcriptional reprogramming orchestrates interactions between tissues continues to be a high priority in regenerative medicine. For example, although initial efforts to generate tissue-specific organoids suggested intrinsic self-organization potential^97^, engineering spatiotemporal cell-cell interactions will likely be needed to improve organoid complexity, function, and translatability^98^. Reprogramming is also likely to influence tubulogenesis and *de novo* establishment of tissue interfaces that occur during heart, lung, kidney, digestive tract, and vascular regeneration, as well as during grafting of engineered tissue and biomaterials^99–101^. Further investigation of how planarian intestinal branches fuse and integrate with the pharynx will provide new insights into these processes.

## Acknowledgments

We thank members of the Forsthoefel lab for valuable insights and critical evaluation of the manuscript. We thank the OMRF Quantitative Analysis Core (Nathan Pezant), Imaging and Histology Core, Center for Biomedical Data Science, and IT/Research Computing Services for invaluable technical assistance. VM was supported by the OMRF John and Mildred Carson Ph.D. Predoctoral Fellowship. DJF was supported by NIH Centers of Biomedical Research Excellence (COBRE) GM103636 (Project 1 to DJF), the Oklahoma Center for Adult Stem Cell Research (a program of TSET) (Project 4340), and the Oklahoma Medical Research Foundation.

## Author Contributions

VMS and DJF designed experiments. VM performed experiments, composed figures, and wrote the initial manuscript draft. DJF and VM conducted bioinformatic analyses, and edited the manuscript.

## Declaration of Interests

The authors declare no competing interests.

## Data and code availability

- Data used to generate the figures are provided in Data S1.
- This paper does not report original code.
- Raw and processed RNA-Sequencing data have been deposited in the NCBI Gene Expression Omnibus (GEO) repository (Accession GSE227155).
- Any additional information required to reanalyze the data reported in this paper is available from the lead contact upon request.

## Experimental model and subject details

Asexual *Schmidtea mediterranea* planarians (clone CIW4)^102^ were cultured in 1X Montjuïc water (1.6 mM NaCl, 1.0 mM CaCl_2_, 1.0 mM MgCl_2_, 0.1 mM KCl, 1.2 mM NaHCO_3_ in deionized water) (double RNAi experiments and Figures 5 and 6), or in Instant Ocean freshwater (0.5 g/L Instant Ocean salts (Spectrum Brands SS3-50) with 0.0167 g/L sodium bicarbonate dissolved in Type I water^103^). Planarians were maintained in plastic Tupperware containers or Petri dishes, incubated in dark at 22°C. Animals were fed calf liver (Skylark, Sprouts Farmers Market, Oklahoma City) homogenized at stored at −80 °C. Wild-type, uninjured animals of similar size, that were not used for prior procedures, were randomly selected and starved 7-10 days before experiments. Animals in experiments were of indeterminate age, since asexual *S. mediterranea* are either very long-lived or immortal^104^.

## Method Details

### Ethics statement

No vertebrate organisms were used in this study.

### *egr* Identification and Nomenclature

Human, mouse, *Drosophila,* and zebrafish Egr protein sequences were used to conduct TBLASTN searches of the Dresden dd_Smed_v6^59^ and Stowers SMED_20160614^57^ transcriptomes, and the Smes_g4 genome assembly^58, 59^. Candidate *S. mediterranea* transcripts predicted to encode three C2H2-type zinc finger domains were identified using NCBI Conserved Domain Finder. Previously undescribed *egr-6, egr-7,* and *egr-8* genes were named in order of discovery according to established nomenclature guidelines^105^.

### Phylogenetic analysis

Predicted triple C2H2-type zinc finger domains in *S. mediterranea* proteins were aligned with zinc finger domains in related transcription factors in other animals in Geneious Prime, using the MUSCLE plug-in with default parameters (PPP algorithm, 0 HMM perturbations and no guide tree permuations). Phylogenetic analysis was conducted using PhyML 3.0 (www.atgc-montpellier.fr/phyml)^106^, with automated smart model selection (Bayesian Information Criterion)^107^ (Q.insect model), BioNJ starting tree, and standard bootstrap analysis (1000 repeats). Resulting Newick tree data were imported into Geneious for dendrogram visualization.

### Cloning and expressed sequence tags

Transcript sequences were identified in the Dresden dd_Smed_v6^59^, Stowers SMED_20160614^57^, and Smed_ESTs3^108^ transcriptomes, and the Smes_g4 genome assembly^58, 59^. Transcript cDNAs were cloned into pBluescript II SK+ or pJC53.2^109^ as previously described^110^, or synthesized by Twist Bioscience (South San Francisco, CA). Full sequences of primers, transcripts, and clones, along with accession numbers, are available in Data S1.

### RNAi

dsRNA synthesis and RNAi experiments were conducted as previously described by mixing 1 or 2 μg of in vitro-synthesized dsRNA with planarian food^110, 111^. For *egr-2* RNAi, animals were fed 4 or 6 times and experiments were initiated 5-7 days after the last feeding. For RNA-Seq, animals were fed 2 μg dsRNA twice per week for 6 times. For double RNAi experiments, animals were fed 5 μg dsRNA of each gene indicated for a total of 10 μg dsRNA twice weekly for 6 times. For *foxF-1* RNAi animals were fed *foxF-1* dsRNA only in the last 1-2 feedings. For *foxA* RNAi animals were fed *foxA* dsRNA in the last 2 feedings. Non-eating planarians were removed if they refused the second dsRNA feeding a day after the regular feeding. For the fluorescent dextran feeding experiment, animals were fed Alexa Fluor 488-conjugated 10,000 MW dextran (Invitrogen D22910) (0.2 mg/ml final concentration) and 2 μg dsRNA the day before imaging.

### In situ hybridization and immunolabeling

Riboprobe synthesis, colorimetric and fluorescent *in situ* hybridizations, and immunolabeling were conducted as previously described^110^. Animals were killed by rocking gently (50 rpm) in 7.5% N-Acetyl-L-Cysteine (NAc, Sigma A7250-100G) for 15 min at RT, followed by fixation in 4% formaldehyde (EMD Millipore FX0410-5) in 1X PBS with Triton X-100 (0.3%) (PBST) for 15 min at RT. Fixed animals were then dehydrated at least O/N in methanol (−30°C) (Sigma A412), rehydrated in PBST and bleached in formamide bleaching solution (4 hours) (5% formamide - Roche 11814320001), and 1.2% hydrogen peroxide (Sigma H1009), diluted in 0.5X SSC (NaCl - Fisher Scientific S271 and sodium citrate - Fisher Scientific S279). Bleached animals were treated with proteinase K (Invitrogen 100005393, 20 μg/ml) in PBSTx (1X PBS - Invitrogen AM9624), 150 mM NaCl, 0.3% Triton X-100 (Fisher Scientific BP151) plus 0.1% SDS (Sigma BP8200) (20-30 min for uninjured animals and 5-20 min for regenerates), followed by 10 min post-fixation. After prehybridization, worms were incubated with Digoxigenin-11-UTP-labeled (Sigma/Roche 11209256910) or Fluorescein-12-UTP-labeled (Sigma/Roche 11427857910) riboprobes (1 ng/μl) for 15-17 hours in hybridization buffer (56°C). After post-hybridization washes, worms were incubated with anti-DIG-AP (Roche/Sigma 11093274910, 1:2000), anti-DIG-POD (Roche/Sigma 11207733910, 1:2000) and/or anti-FITC-HRP (Roche/Sigma 11426346910, 1:2000) for 15-17 hours in TNTx (Tris-HCl pH 8.0 - Invitrogen 15567-027, 150 mM NaCl, 0.3% Triton X-100) with 5% horse serum (Sigma H1138-500ML) and 0.5% Roche Western Blocking Reagent (Roche/Sigma 11921673001). Animals were then developed with colorimetric substrate (NBT/BCIP) or by tyramide signal amplification (fluor-tyramide at 1:1000-1:2000, 10 min, RT) (tyramides synthesized exactly as described^62^). For double FISH, HRP was inactivated by incubating with 100 mM NaN_3_ (Fisher Scientific S227I) for 45 minutes. DAPI labeling (1 μg/ml) (Sigma D9542) was performed either with antibody incubation, or during post-TSA washes.

For muscle immunolabeling, samples were incubated with mouse 6G10 anti-muscle (Developmental Studies Hybridoma Bank, 6G10-2C7) (1:250)^112^ in IF block (1X PBS, 0.45% fish gelatin (Sigma G7765-250ML), 0.6% IgG-free BSA (Jackson ImmunoResearch 001-000-162), 0.3% Triton X-100) O/N at 4°C. Samples were washed 6 times (10 min, RT) with PBSTx after antibody incubation. Samples were then incubated with goat anti-mouse-488 (1:250) (Jackson ImmunoResearch, 115-545-146) O/N at 4°C. Samples were washed 6 times in PBSTx.

Samples were mounted in Vectashield (Vector Labs H-1000) or Fluoromount G (Southern Biotech 0100-01).

### RNA extraction, library preparation, and RNA sequencing

Anterior tissue from ∼10 tail regenerates per biological replicate was homogenized in Trizol (Thermo Fisher Scientific 15596026) with a motorized Kontes pestle grinder. RNA was extracted with two chloroform extractions and high-salt precipitation buffer according as per the Trizol protocol. Precipitated RNA solutions were transferred to Zymo RNA columns (Zymo RNA Clean & Concentrator 5 kit, R1013) for DNAse treatment and purification as per manufacturer’s protocol. RNA samples were submitted to GENEWIZ (South Plainfield, NJ) for library generation using NEB NEXT ULTRA library prep (New England Biolabs) and RNA sequencing with standard Illumina adapters. Paired-end (2 x 150 bp) sequence was generated on an Illumina HiSeq 4000 instrument; 19 M–26 M reads were generated for each replicate.

### Read mapping

Quality control and read mapping to unique transcripts in dd_Smed_v6^113^ were conducted with FastQC (v0.11.5)^114^, BBDuk (v35.66) (https://sourceforge.net/projects/bbmap/), and Bowtie2 (v2.3.1)^115^. BBDuk (v36.99) settings for paired end reads: k=13 ktrim=r mink=11 qtrim=rl trimq=10 minlength=35 tbo tpe. Bowtie2 (v2.3.1) for paired end reads was used for mapping, with “-a” multi-mapping and “–local” soft-clipping allowed. For read summarization, the “featureCounts” utility in the Subread package (v1.6.3)^116^ was used with a custom “.SAF” file and options “-p-M-O-F SAF” to include multi-mapping and multi-overlapping reads.

### Differential expression analysis

Read counts matrices were imported into R v4.1.0^117^ and analyzed in edgeR v3.34.1^118^. All transcripts with counts per million (CPM) < 1 (lowly expressed transcripts) in four samples were excluded from further analysis. Next, after recalculation of library sizes, samples were normalized using trimmed mean of M-values (TMM) method, followed by calculation of common, trended, and tagwise dispersions. Then, differentially expressed transcripts were identified using the generalized linear model (GLM) likelihood ratio test. Expression changes were considered significant if the false discovery rate-adjusted *P* value (“FDR”) was <0.05. Expression data are provided in Tables S1-S4.

For cross-referencing, *egr-2(RNAi)* expression data were merged in R with lists of transcripts associated with the early injury response^37^, polarity^37, 66, 67^, muscle subtypes^64^, and specific cell subpopulations^70^. To increase the likelihood of detecting changes in specific cell types and tissues, we limited analysis to the most (top 25 percent) enriched transcripts in each cell type (based on log-fold-enrichment scores, 4,078 transcripts), to limit the representation of transcripts expressed by multiple cell types^70^.

### Single cell data analysis

Single cell gene expression matrices for unirradiated samples (Days 0, 1, 2, 4, 7, 10, and 14, GSM4404045-GSM4404051) were downloaded from the NCBI Gene Expression Omnibus (GSE146685) and processed in Seurat 4.1.1^119^ in R 4.1.2 on the Oklahoma Medical Research Foundation compute cluster (platform: x86_64-pc-linux-gnu running under Ubuntu 20.04.4 LTS). Only features expressed by at least three cells, and only cells expressing at least 75 features (nGenes/nFeatures) (e.g., SMED20140614 transcripts) and at least 500 total transcripts (nUMI/nCounts) were retained for analysis^65^. In addition, cells with more than five percent mitochondrial RNA (SMED30012212, SMED30008414, SMED30019959, SMED30026531, SMED30000702, SMED30003686, SMED30010375, SMED30001799)^67^ were removed. Data were normalized using SCTransform^65, 120, 121^ with the glmGamPoi package and integrated with nfeatures=3000 (https://satijalab.org/seurat/articles/integration_introduction.html, compiled 2022-01-11). Default parameters were used for all steps, except PCA dimensionality reduction was performed with npcs=200, and UMAP embedding was optimized by testing 25, 50, 75, 100, 150, and 200 dimensions; dims=100 yielded 35 clusters/subclusters analogous to previously identified planarian cell types and states^65, 70, 122^. For co-expression, all cells with non-zero *egr-2* expression (SMED30035904) were subsetted for generation of UMAP plots of transcript co-expression in *egr-2-*positive cells.

### qRT-PCR

Total RNA was extracted from 10 trunk fragments per biological replicate using the standard Trizol RNA extraction protocol with high salt precipitation and clean-up on Zymo RNA columns (Zymo R1013/R1017). RNA (1 μg) was reverse transcribed using the iScript cDNA Synthesis kit (BioRad 1708890). mRNA expression was detected using the Fast Start Essential Green DNA master mix (Roche 06924204001) on a Roche LightCycler 96. Relative mRNA expression levels were normalized to the geometric mean of endogenous controls *ef-2* and *gapdh* using the Livak ΔΔCt method^123^.

### Image Collection and Quantification

Images of live animals, live regenerates, and colorimetric *in situ* samples were collected on a Zeiss Stemi 508 with an Axiocam 105 color camera (ZEN 2.3 Blue). Epifluorescent images were collected on a Zeiss AxioObserver.Z1 with Excelitas X-Cite 120 LED Boost illumination and ZEN 2.3 Blue (v2.3.64.0). Live dextran-fed animals were imaged with a Leica M205 FCA (LAS X v5.0.2.24429). Confocal images (Figures 2A, 2B, 2D, 2E, 4E, 5E, 5G, and 5I) were collected on a Zeiss LSM 710 laser scanning microscope with 10X Plan NeoFluar, 20X Plan Apo, or 40X C-Apo objectives (ZEN version 11.0.3.190, 2012-SP2).

For anterior intestine phenotypes, fragments were scored as having a “split” phenotype if the anteriormost intestinal branches failed to fuse at the midline for at least one third the total length of the anterior intestine. Fragments were scored as having “gaps” if the intestine had fused at the anterior but one or more smaller regions, not labeled by intestine-specific riboprobes but surrounded by intestinal tissue, were present in a midline region approximately one third the total intestine/animal width. Gap size was scored in ImageJ by tracing all gaps per fragment, calculating the sum of these areas, and normalizing to total fragment area. Fragments scored as “anterior fusion only” only had midline intestinal tissue at the anteriormost extent of the intestine, while in “no fusion” animals, intestine was absent across the entire anterior (e.g. ∼50% fragment length) midline.

For quantification of animal area, blastema area, brain and pharynx size, regions were traced in ImageJ^124^, measured, and organ/region-to-body size ratios were calculated. For quantification of colorimetric *sFRP-1* expression, area covered by mRNA-positive cells was manually traced, measured, and normalized to total animal area. For quantification of colorimetric *slit-1* expression, images were converted to 8-bit, then *slit-1-*positive regions in the anterior 33% of each fragment were delineated with the Adjust Threshold tool. After automated tracing with the Wand Tool, area was measured and normalized to total animal area.

For *mp-1+* cell quantification, animals or fragments were imaged ventrally using Zeiss 710 confocal microscope. For intact animals, a rectangle was drawn at the midline between the posterior gut branches. The width of the box was set as the 10% of the posterior gut length (distance from posterior pharynx to posterior gut tip) and the length of the box was set as 20% of the posterior gut length. For tail regenerates, the width of the box was 50% of pharynx width and the length extended from the tip of the anterior intestine to the pharynx attachment site. Five maximum intensity projections of three optical sections per projection (0.471 μm step size) were generated per animal/fragment, skipping two sections between projections to avoid counting cells twice. Imaging was initiated at approximately half the distance between the ventral epidermis and the intestine, and progressed dorsally. *mp-1+* cells were counted manually in each max projection using the multi-point tool in FIJI. m*p-1+* cells per animal were quantified by dividing the sum of *mp-1+* cells across the five maximum intensity projections over the total volume of the imaged z-stacks (5.652μm^3^).

For *slit-1* fluorescence in the presumptive pharynx region, fragments were imaged with identical exposure times. Then, in ImageJ, a box with 15% animal length was drawn across the full animal width (based on DAPI fluorescence) and positioned over the central region of highest *slit-1* intensity (∼20-35% of animal length for 2 dpa fragments, and ∼35-50% for 3 dpa fragments) (see Fig. S7C). The Plot Profile tool was then used to quantify vertically averaged pixel intensity across the x-axis within this box. Pixel intensities were also calculated in a second background box with 15% animal length and 10% animal width, positioned at the posterior midline. Profile data were exported to spreadsheets, where x-axis positions in box 1 were normalized to a maximum width of 1. For each fragment, average background fluorescence in box 2 was subtracted from fluorescence intensity at each position across 0.4-0.6 animal width (the region of pharynx-associated *slit-1* expression), then averaged.

For displacement of *piwi-1+* cells at the presumptive pharyngeal cavity, fragments were imaged with identical exposure times. Then, in ImageJ, a box with a mediolateral width equal to 20% of fragment length was drawn at the midline, covering 20% to 50% length from anterior to posterior (see Fig S7D). The Plot Profile tool was used to quantify vertically averaged pixel intensity across the x-axis within this box. Profile data were normalized so that the highest peak intensity (to either the left or right of the pharyngeal cavity) was 1; then, the lowest midline intensity between the two peaks was subtracted from one and multiplied by 100 to calculate the percentage decrease in midline *piwi-1* signal.

### Quantification and Statistical Analysis

All statistical tests were performed in Prism 9 (GraphPad Prism Software, San Diego, CA). The number of values (n) are indicated by the number of data points in figure plots, along with standard deviation error bars and *P* values. n values and source data are provided for each experiment in Data S1.

Replicate information: At least two independent FISH or immunolabeling experiments with at least five animals per condition were performed for qualitative characterization of gut, muscle, or polarity marker phenotypes. Colorimetric in situ hybridization time course on all *egr* genes was conducted with 5-10 fragments per condition. Number of animals or fragments for all other experiments are provided in Data S1.

Two way ANOVA was used only when time’s influence on an RNAi phenotype (Figure 5H and Figure S4B) or knockdown efficiency (Figure S3A) was evaluated. One way ANOVA was used to compare multiple knockdown combinations. For intestine gap size comparisons, outliers (highlighted in Data S1) were removed using Grubbs’ test; otherwise, all animals/fragments analyzed are included. Chi-square or Fisher’s exact test was used to compare number of fragments displaying phenotypes. Welch’s *t* test was used for all other pairwise comparisons.

Statistical analysis information for RNA-Seq experiments are described in the corresponding Methods sections.

---

**Figure S1.**
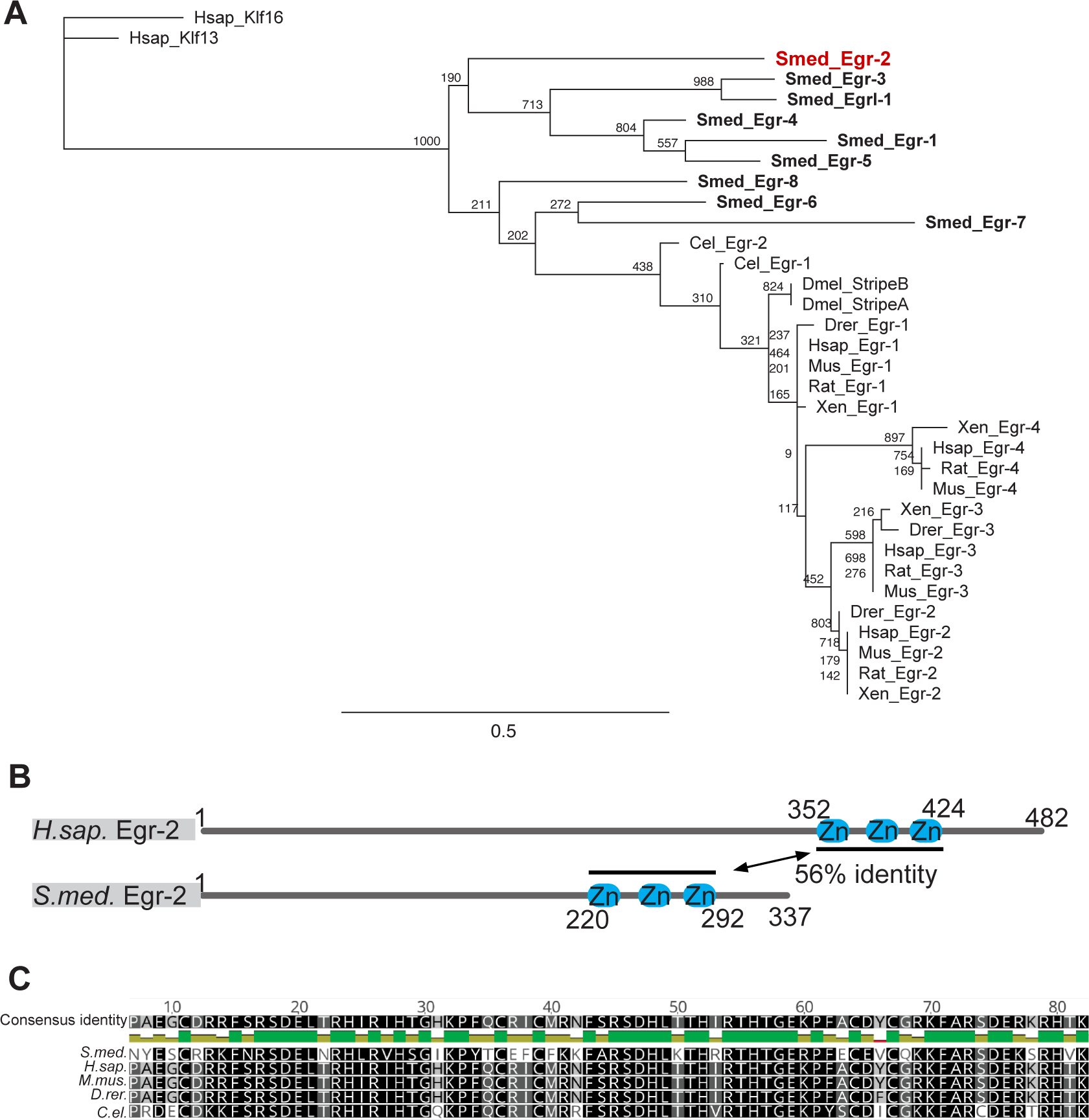
Evolutionary conservation of Egr transcription factors. **(A)** Phylogenetic analysis of C2H2-Zn-finger domains encoded by planarian (“Smed”) *egr* homologs and related genes in other species. Planarian Egr proteins group with other Egr proteins, and not closely-related Krüppel-like factor (Klf) proteins. Maximum-likelihood phylogeny with 1000 bootstrap replicates. Branch support is indicated. **(B)** Domain alignment of C2H2-Zn-finger domains in human and *S. mediterranea* Egr-2. **(C)** Alignment of Egr-2 C2H2-Zn-finger domains from multiple species showing amino acid similarity and identity.

**Figure S2.**
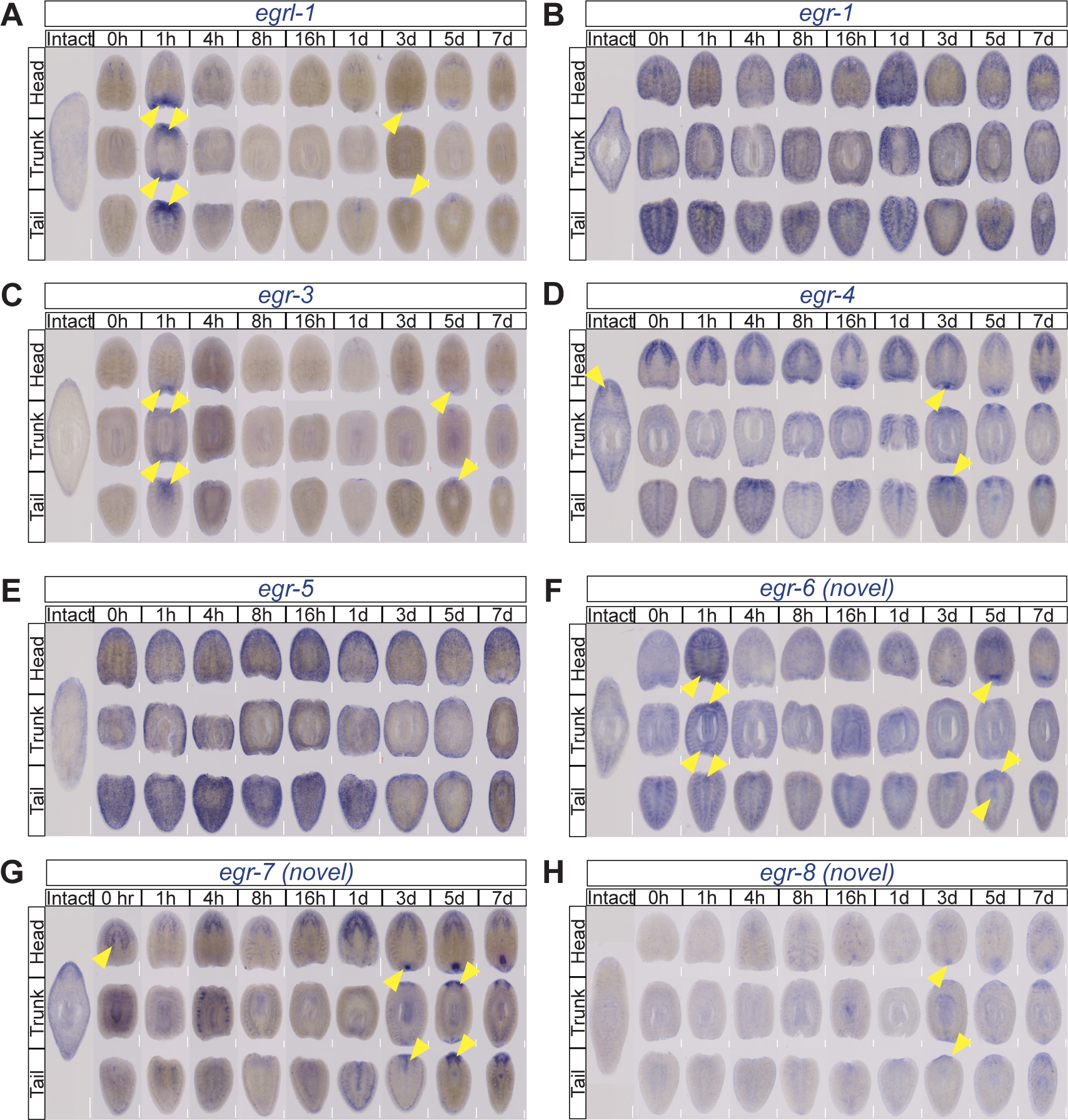
Expression of *egr* transcripts during regeneration. **(A-H)** Colorimetric ISH of each *egr* during a 7 dpa regeneration time course. Yellow arrows, regions of expression or upregulation during regeneration. Dorsal view, anterior up. Scale bar: 200μm. **(A)** *egrl-1* was transiently upregulated in wound proximal regions by 1 hpa. Modest expression was also observed at the amputation plane in head and tail fragments. **(B)** *egr-1* was expressed in epidermal and parenchymal cells and is not wound-induced. **(C)** *egr-3* was transiently upregulated in wound proximal regions by 1 hpa and in the blastema region by 3-5 dpa. **(D)** *egr-4* was expressed in the central nervous system and the regenerating pharynx and nervous system. **(E)** *egr-5* was expressed in epidermal progenitor cells and epidermal progeny and was not wound-induced. **(F)** *egr-6* (novel) was expressed in parenchymal cells and the regenerating pharynx and blastema. **(G)** *egr-7* (novel) was expressed in the pre-existing and regenerating nervous system and regenerating pharynx. *(A)−8* (novel) was expressed in the regenerating pharynx and blastema.

**Figure S3.**
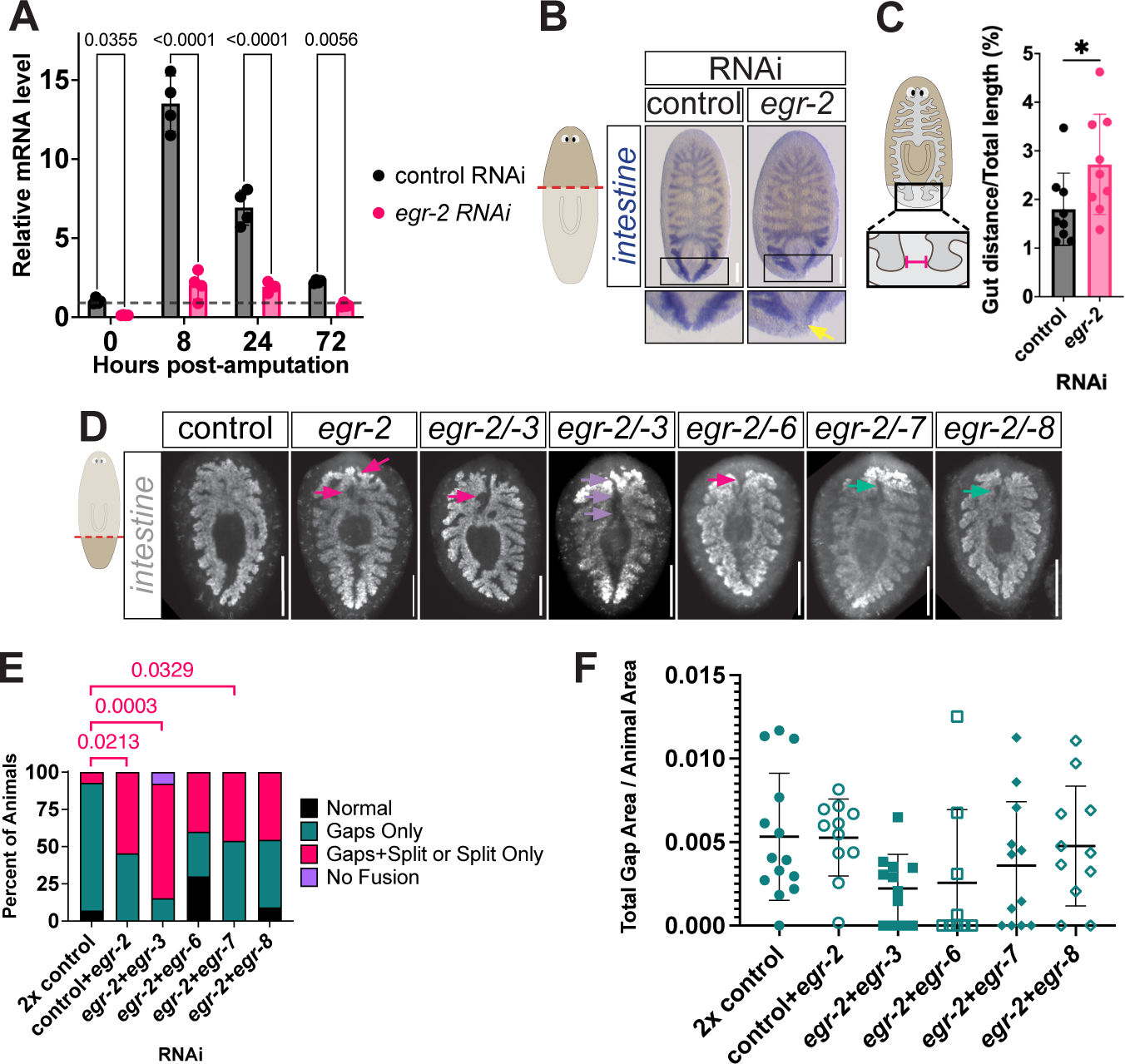
RNAi screen and phenotype scoring for *egr* genes. **(A)** qPCR analysis of mRNA levels in control and *egr-2(RNAi)* fragments. Dashed line, relative mRNA level of 1. Error bars, mean ±SD. Dots, n=4 biological replicates per condition per time point, 10 trunk fragments per biological replicate. Two-way ANOVA with Benjamini, Krieger, and Yekutieli correction for multiple comparisons (control and *egr-2(RNAi)* comparisons only at each time point). *q* values shown. **(B)** Colorimetric ISH (*apob-2* riboprobe) labeling the intestine in head regenerates (6 dpa). Arrow, wider gap between regenerating tail branches. Dorsal view, anterior up. Scale bar, 200μm. **(C)** Quantification of distance between gut branches from (B) in the posterior of head regenerates relative to total animal length. Error bars, mean ±SD. Points, individual animals. Welch’s *t* test (\**P*=0.0462). **(D)** Fluorescent ISH with intestine riboprobe pool on 6 dpa tail fragments after *egr-2* RNAi in combination with other *egr* homologs. Magenta arrows: split anterior intestinal branches. Teal arrows: intestinal gaps. Purple arrows: no fusion. Dorsal view, anterior up. Scale bar, 200μm. **(E)** Phenotype scoring in (F). Fisher’s exact test, *P* values shown for fragments with split or split plus gaps phenotypes. Number of fragments with gaps only or no fusion were not significantly different. **(F)** Normalized total gap area. Points indicate total gap area per animal divided by total animal area. Error bars, mean ±SD. Ordinary one-way ANOVA with Sídâk correction for multiple comparisons (no significant differences).

**Figure S4.**
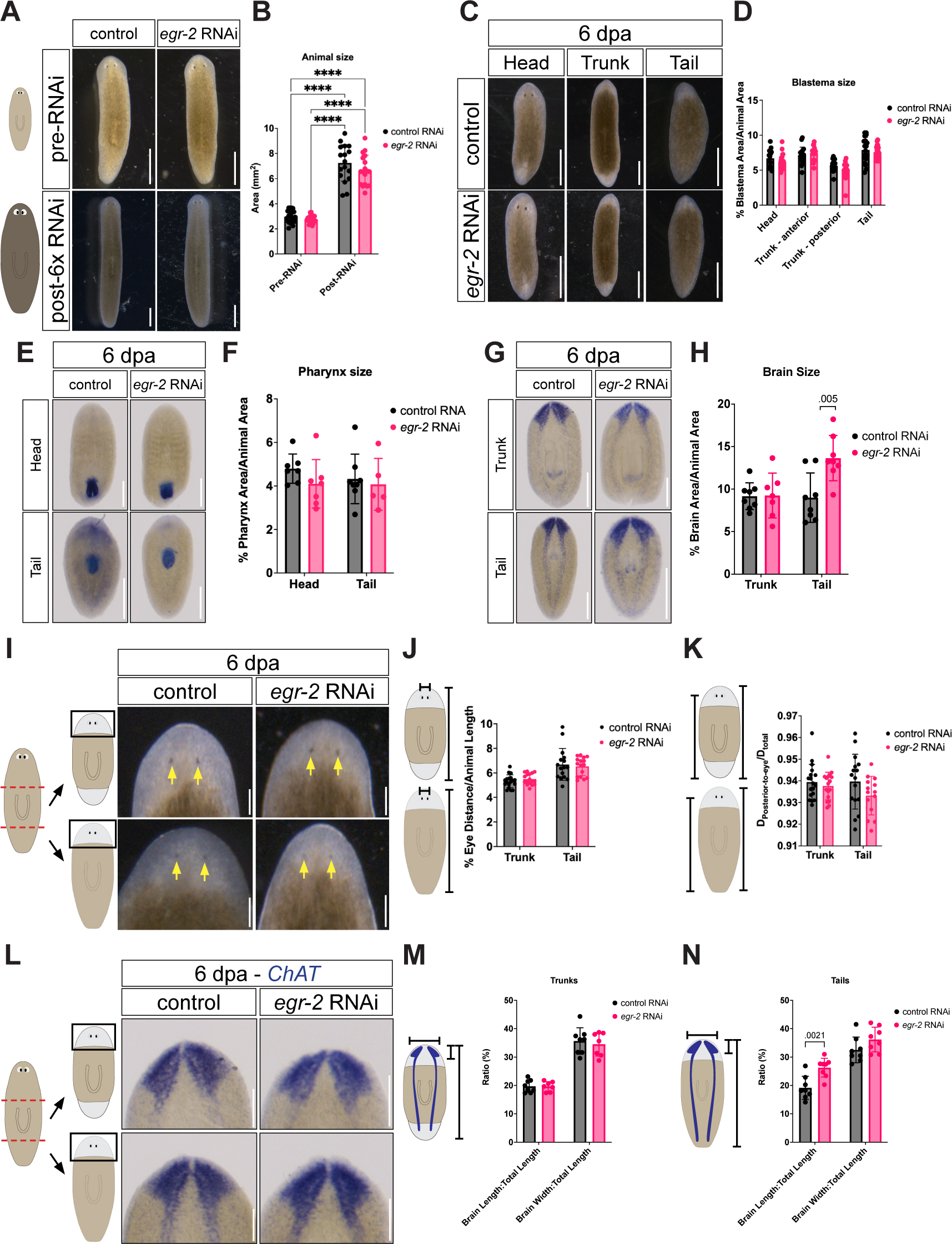
*egr-2* RNAi does not affect growth or regeneration of blastema, brain, eyes, or scaling. **(A)** Uninjured animals before and after six dsRNA feedings. Scale bars, 1 mm. **(B)** Animal area before and after 6 RNAi feedings. *egr-2* RNAi did not affect animal growth. Two-way ANOVA with Tukey’s multiple comparisons test, \*\*\*\**P*<.0001. **(C)** Six day regenerates. Scale bars, 1 mm. **(D)** Normalized blastema size at anterior and posterior wounds in six day regenerates. *egr-2* RNAi did not affect blastema size. Welch’s *t* test (control vs. RNAi for each fragment type), no significant differences. **(E)** Colorimetric ISH with *laminin* riboprobe labeling pharynx in 6 dpa head (top) and tail (bottom) regenerates. Scale bars, 200 μm. **(F)** Normalized pharynx size in regenerates. *egr-2* RNAi did not affect pharynx size. Welch’s *t* test (control vs. RNAi for each fragment type), no significant differences. **(G)** Colorimetric ISH with *ChAT* riboprobe labeling brain in 6 dpa trunk (top) and tail (bottom) regenerates. Scale bars, 200 μm. **(H)** Normalized brain size in regenerates. *egr-2* RNAi did not affect brain size in trunks. *egr-2(RNAi)* tail fragments were smaller than controls, causing brains to be proportionally larger (see Data S1). Welch’s *t* test (control vs. RNAi for each fragment type), no significant differences in trunks; *P* value shown for tail comparison. **(I)** Regenerating eyes (arrows) in 6 dpa trunk (top) and tail (bottom) regenerates. Scale bars, 200μm. **(J)** Distance between eyes normalized to animal length. *egr-2* RNAi did not affect inter-eye distance. Welch’s *t* test (control vs. RNAi for each fragment type), no significant differences. **(K)** Distance between eyes and posterior tip normalized to animal length. *egr-2* RNAi did not affect eye positioning. Welch’s *t* test (control vs. RNAi for each fragment type), no significant differences. **(L)** Colorimetric ISH with *ChAT* riboprobe labeling brain in 6 dpa tail regenerates. Scale bars, 100 μm. **(M)** Brain length and width normalized to trunk fragment length. *egr-2* RNAi did not affect brain proportions. Welch’s *t* test (control vs. RNAi), no significant differences. **(N)** Brain length and width normalized to tail fragment length. *egr-2* RNAi did not affect brain width, but brain length was proportionally larger since *egr-2(RNAi)* fragments were smaller than controls (see Data S1). Welch’s *t* test (control vs. RNAi), *P* value shown for brain length; width was not significantly different. In (B, D, F, H, J, K, and M), points represent values from individual animals. Error bars = mean ±SD.

**Figure S5.**
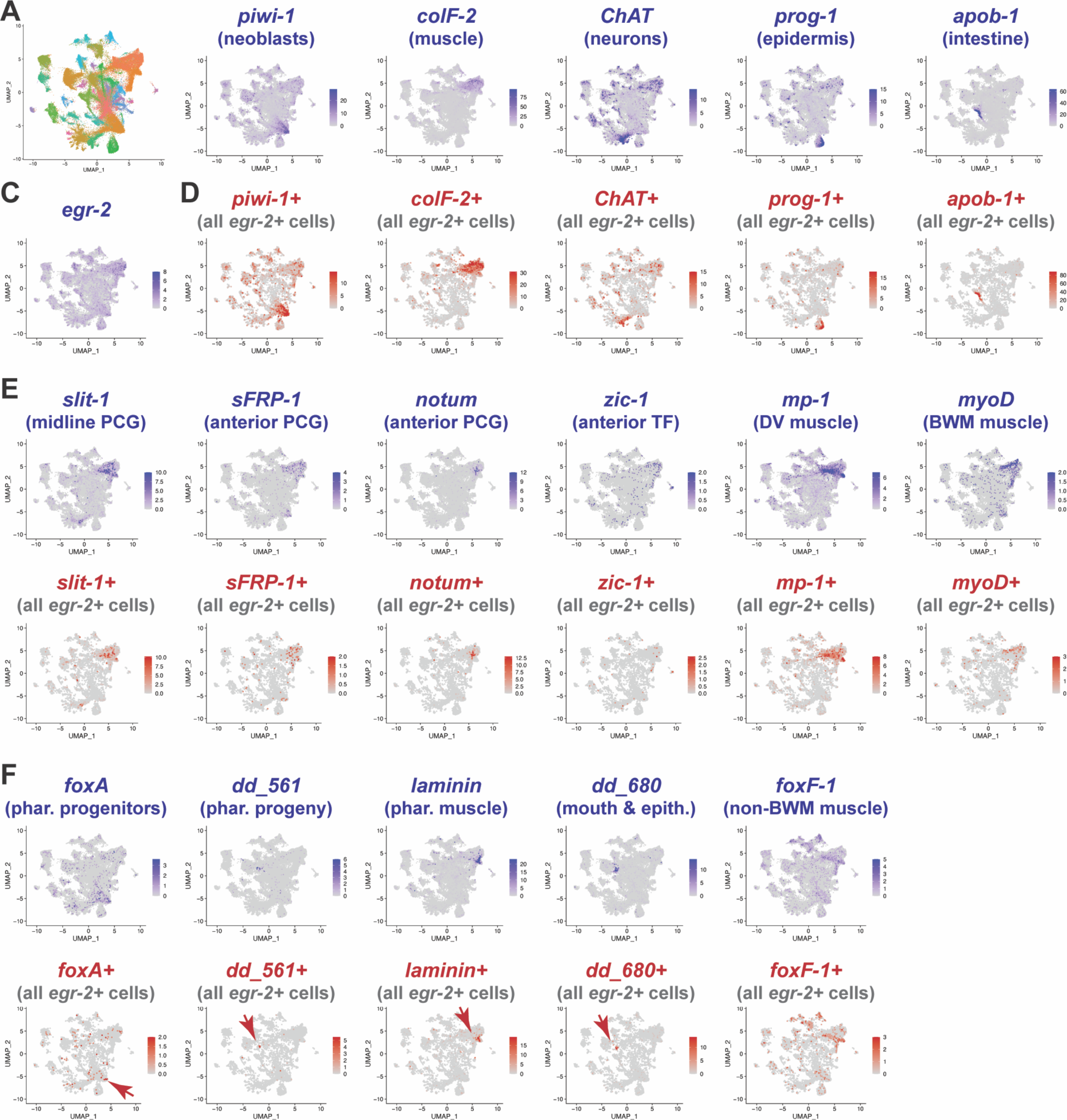
*egr-2* mRNA is expressed in muscle and pharynx-related cell types. **(A)** UMAP plot showing 35 cell type/subtype clusters in re-analyzed tissue fragment data from unirradiated samples in Benham-Pyle et al., 2021. **(B)** UMAP feature plots showing expression (blue) of cell type-enriched transcripts. **(C)** UMAP feature plot showing *egr-2* expression in multiple tissues. **(D)** UMAP feature plots of only *egr-2-*positive (gray) cells showing co-expression of cell type-enriched transcripts (red). **(E)** UMAP feature plots of polarity-associated transcripts (PCG, position control gene) and muscle subtype markers in all cells (top, blue) and in *egr-2-*positive cells (bottom, red). **(F)** UMAP feature plots of pharynx-associated transcripts in all cells (top, blue), and in *egr-2-* positive cells (bottom, red). Color bars, non-normalized RNA counts for each transcript. For all-cell plots (blue) except *piwi-1* and *colF-2*, *egr-2*, and *foxF-1*, the top 1% of cells expressing the highest levels of each transcript were not shown to increase contrast. For *egr-2-*positive (red) cell plots, the top 5% (*foxA, sFRP-1*) or 1% (all others) of cells were excluded.

**Figure S6.**
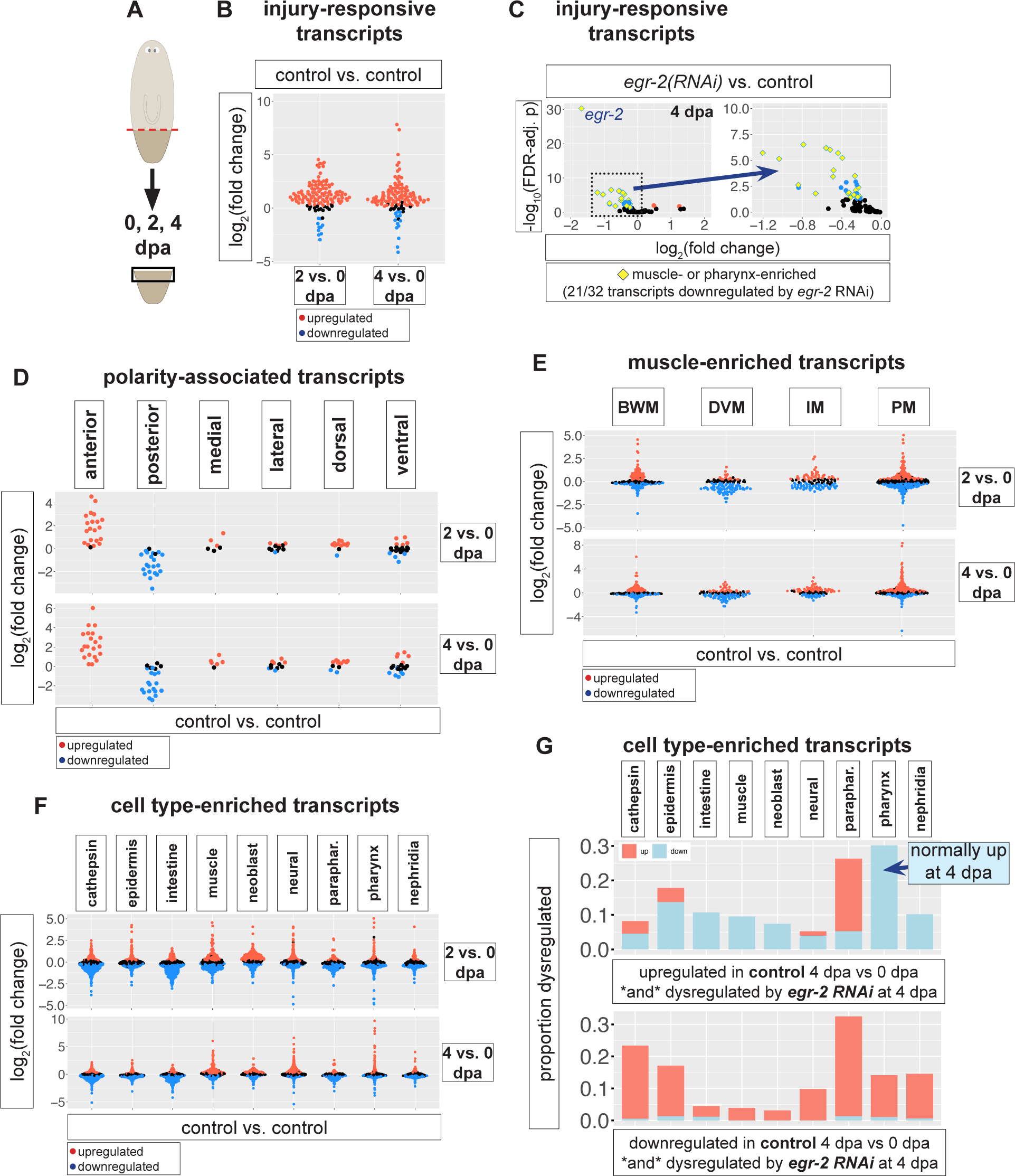
Injury-responsive transcript expression in control and *egr-2(RNAi)* tissue. **(A)** Schematic of tissue isolation strategy. **(B)** Differential expression of injury-responsive transcripts at 2 and 4 dpa vs. 0 dpa in control wound-proximal tissue. **(C)** Volcano plots showing 21/32 injury-responsive transcripts downregulated by *egr-2* RNAi at 4 dpa are also normally enriched in muscle or pharynx in uninjured animals (yellow diamonds). **(D)** Expression of polarity-associated transcripts at 2 and 4 dpa vs. 0 dpa in control tissue. **(E)** Expression of muscle-enriched transcripts at 2 and 4 dpa vs. 0 dpa in control tissue. BWM, body wall muscle. DVM, dorsoventral muscle. IM, intestinal muscle. PM, pharynx muscle. **(F)** Expression of cell type-enriched transcripts at 2 and 4 dpa vs. 0 dpa in control tissue. **(G)** Percentage of cell type-enriched and normally injury-responsive transcripts dysregulated by *egr-2* RNAi at 4 dpa. In B-F, significantly up-or down-regulated transcripts (FDR-adjusted *p* value < 0.05) are shown in red or blue, respectively.

**Figure S7.**
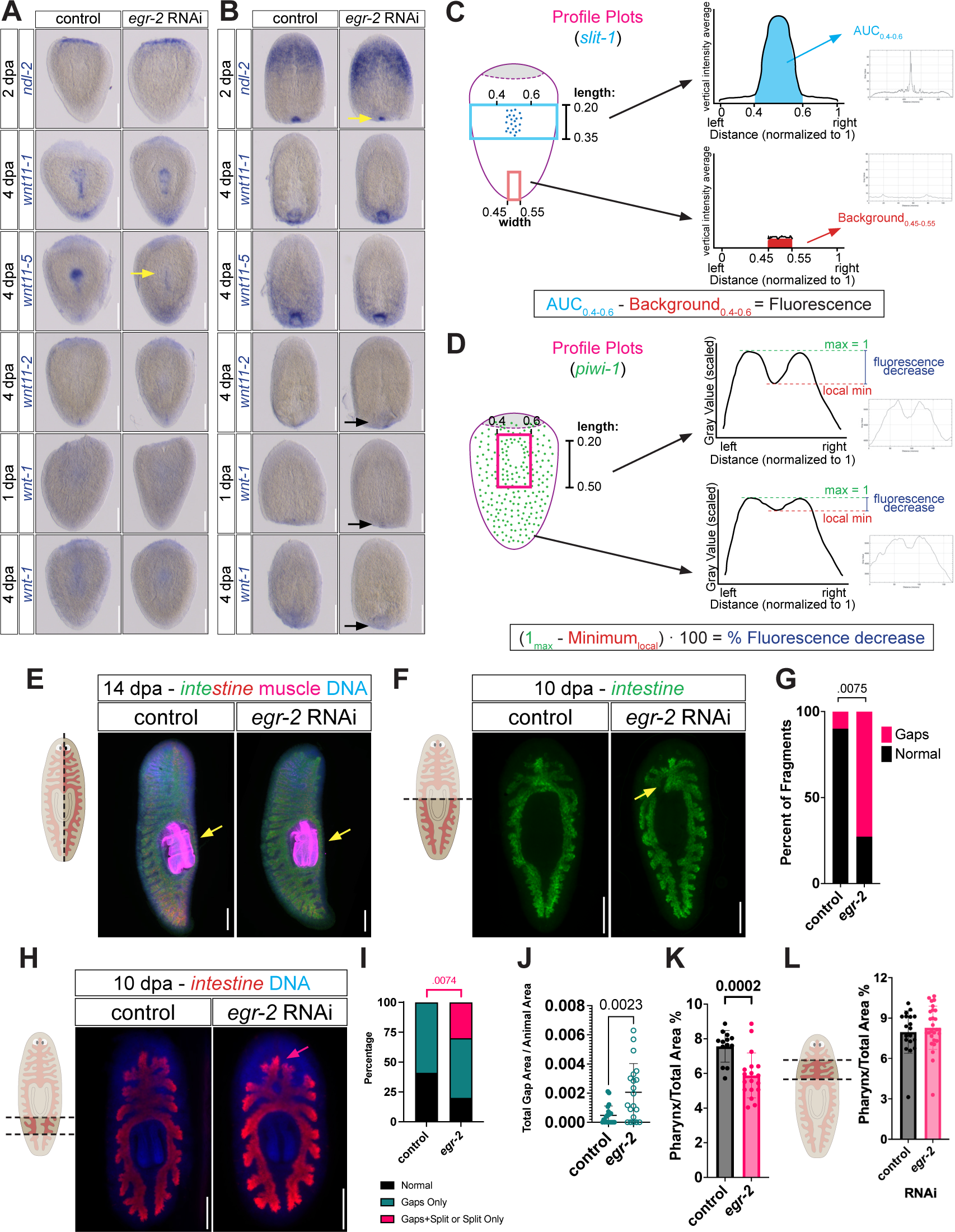
Polarity gene expression, midline fluorescence quantification strategies, and *egr-2* dependence of anterior intestine remodeling when pharynx regeneration is required. **(A)** Expression of anterior (*ndl-2),* pharynx/posterior (*wnt11-1, wnt11-5*), and posterior (*wnt11-2, wnt-1*) polarity cues (colorimetric ISH) in tail fragments. Yellow arrow, decreased *wnt11-5* expression. Dorsal view, anterior up. Scale bars, 200 μm. **(B)** Expression of polarity cues in head fragments. Yellow arrow, decreased *ndl-2* expression. Black arrows, re-expression of posterior cues. Dorsal view, anterior up. Scale bars, 200 μm. **(C)** Strategy for quantification of *slit-1* expression (detected by fluorescent ISH) at the ventral midline/presumptive pharynx regions. Detailed description in Methods. **(D)** Strategy for quantification of *piwi-1* expression (detected by fluorescent ISH) at the ventral midline/presumptive pharyngeal cavity. Detailed description in methods. **(E)** Fluorescent ISH with intestine riboprobe pool (green/magenta), muscle IF (anti-6G10, magenta only), and DAPI (blue) in 14 dpa sagittally amputated fragments. Arrows, regenerated peri-pharyngeal intestinal branch. Dorsal view, anterior up. Scale bar, 200 μm. **(F)** Fluorescent ISH with intestine riboprobe pool in 10 dpa posterior fragments amputated transversely through the pharynx. Arrow, gap at pharynx attachment site. Dorsal view, anterior up. Scale bar, 200 μm. **(G)** Quantification of pharynx attachment site gaps in animals from **(F).** Control n=12, *egr-2(RNAi)* n=9. Fisher’s exact test, *P* value shown. **(H)** Normalized pharynx size in thin posterior transverse fragments. Points, values for individual fragments. Welch’s *t* test, *P* value shown. **(I)** Normalized pharynx size in thin anterior transverse fragments. Points, values for individual fragments. Welch’s *t* test, not significant. **(J)** Fluorescent ISH with intestine riboprobe pool in 10 dpa thin posterior fragments. Arrow, split intestine. Dorsal view, anterior up. Scale bar, 200 μm. **(K)** Quantification of phenotypes in **(I).** Fisher’s exact test, *P* value shown; there was no significant difference in the number of animals with gaps. **(L)** Total gap area normalized to fragment area. Welch’s *t* test, *P* value shown.

## Notes

### Competing Interest Statement

The authors have declared no competing interest.

## References

1. Tanaka, E.M., and Reddien, P.W. (2011). The cellular basis for animal regeneration. Dev. Cell 21, 172–185. 10.1016/j.devcel.2011.06.016.

2. Sánchez Alvarado, A., and Yamanaka, S. (2014). Rethinking differentiation: stem cells, regeneration, and plasticity. Cell 157, 110–119. 10.1016/j.cell.2014.02.041.

3. Ricci, L., and Srivastava, M. (2018). Wound-induced cell proliferation during animal regeneration. Wiley Interdiscip Rev Dev Biol 7, e321. 10.1002/wdev.321.

4. Niethammer, P. (2016). The early wound signals. Curr. Opin. Genet. Dev. 40, 17–22. 10.1016/j.gde.2016.05.001.

5. Goldman, J.A., and Poss, K.D. (2020). Gene regulatory programmes of tissue regeneration. Nat. Rev. Genet. 21, 511–525. 10.1038/s41576-020-0239-7.

6. Duncan, E.M., and Sánchez Alvarado, A. (2019). Regulation of Genomic Output and (Pluri)potency in Regeneration. Annu. Rev. Genet. 53, 327–346. 10.1146/annurev-genet-112618-043733.

7. Srivastava, M. (2021). Beyond Casual Resemblance: Rigorous Frameworks for Comparing Regeneration Across Species. Annu. Rev. Cell. Dev. Biol. 37, 415–440. 10.1146/annurev-cellbio-120319-114716.

8. Thornton, C.S. (1957). The effect of apical cap removal on limb regeneration in Amblystoma larvae. J Exp Zool 134, 357–381. 10.1002/jez.1401340209.

9. Godwin, J.W., Pinto, A.R., and Rosenthal, N.A. (2013). Macrophages are required for adult salamander limb regeneration. Proc Natl Acad Sci U S A 110, 9415–9420. 10.1073/pnas.1300290110.

10. Petrie, T.A., Strand, N.S., Yang, C.T., Rabinowitz, J.S., and Moon, R.T. (2014). Macrophages modulate adult zebrafish tail fin regeneration. Development 141, 2581–2591. 10.1242/dev.098459.

11. Simkin, J., Gawriluk, T.R., Gensel, J.C., and Seifert, A.W. (2017). Macrophages are necessary for epimorphic regeneration in African spiny mice. Elife 6. 10.7554/eLife.24623.

12. Sun, J., Peterson, E.A., Wang, A.Z., Ou, J., Smith, K.E., Poss, K.D., and Wang, J. (2022). *hapln1* Defines an Epicardial Cell Subpopulation Required for Cardiomyocyte Expansion During Heart Morphogenesis and Regeneration. Circulation 146, 48–63. 10.1161/CIRCULATIONAHA.121.055468.

13. Biswas, L., Chen, J., De Angelis, J., Singh, A., Owen-Woods, C., Ding, Z., Pujol, J.M., Kumar, N., Zeng, F., Ramasamy, S.K., and Kusumbe, A.P. (2023). Lymphatic vessels in bone support regeneration after injury. Cell 186, 382–397 e324. 10.1016/j.cell.2022.12.031.

14. Newmark, P.A., and Sánchez Alvarado, A. (2002). Not your father’s planarian: a classic model enters the era of functional genomics. Nat. Rev. Genet. 3, 210–219. 10.1038/nrg759.

15. Reddien, P.W., and Sánchez Alvarado, A. (2004). Fundamentals of planarian regeneration. Annu. Rev. Cell Dev. Biol. 20, 725–757. 10.1146/annurev.cellbio.20.010403.095114.

16. Gurley, K.A., Elliott, S.A., Simakov, O., Schmidt, H.A., Holstein, T.W., and Sánchez Alvarado, A. (2010). Expression of secreted Wnt pathway components reveals unexpected complexity of the planarian amputation response. Dev. Biol. 347, 24–39. 10.1016/j.ydbio.2010.08.007.

17. Forsthoefel, D.J., Park, A.E., and Newmark, P.A. (2011). Stem cell-based growth, regeneration, and remodeling of the planarian intestine. Dev. Biol. 356, 445–459. 10.1016/j.ydbio.2011.05.669.

18. Umesono, Y., Tasaki, J., Nishimura, Y., Hrouda, M., Kawaguchi, E., Yazawa, S., Nishimura, O., Hosoda, K., Inoue, T., and Agata, K. (2013). The molecular logic for planarian regeneration along the anterior-posterior axis. Nature 500, 73–76. 10.1038/nature12359.

19. Asai, E. (1991). Regeneration of the pharynx in a freshwater planarian: an electronmicroscopic study with special reference to the formation of the pharyngeal cavity and pharyngeal lumen. Zool. Sci. 8, 775–784. 10.34425/zs000897.

20. Bueno, D., Espinosa, L., Baguñà, J., and Romero, R. (1997). Planarian pharynx regeneration in regenerating tail fragments monitored with cell-specific monoclonal antibodies. Dev. Genes Evol. 206, 425–434. 10.1007/s004270050072.

21. Kobayashi, C., Watanabe, K., and Agata, K. (1999). The process of pharynx regeneration in planarians. Dev. Biol. 211, 27–38. 10.1006/dbio.1999.9291.

22. Adler, C.E., Seidel, C.W., McKinney, S.A., and Sánchez Alvarado, A. (2014). Selective amputation of the pharynx identifies a FoxA-dependent regeneration program in planaria. Elife 3, e02238. 10.7554/eLife.02238.

23. Cebrià, F., Guo, T., Jopek, J., and Newmark, P.A. (2007). Regeneration and maintenance of the planarian midline is regulated by a *slit* orthologue. Dev. Biol. 307, 394–406. 10.1016/j.ydbio.2007.05.006.

24. Gurley, K.A., Rink, J.C., and Sánchez Alvarado, A. (2008). β-catenin defines head versus tail identity during planarian regeneration and homeostasis. Science 319, 323–327. 10.1126/science.1150029.

25. Petersen, C.P., and Reddien, P.W. (2008). *Smed-βcatenin-1* is required for anteroposterior blastema polarity in planarian regeneration. Science 319, 327–330. 10.1126/science.1149943.

26. Adell, T., Saló, E., Boutros, M., and Bartscherer, K. (2009). Smed-Evi/Wntless is required for β-catenin-dependent and-independent processes during planarian regeneration. Development 136, 905–910. 10.1242/dev.033761.

27. Yazawa, S., Umesono, Y., Hayashi, T., Tarui, H., and Agata, K. (2009). Planarian Hedgehog/Patched establishes anterior-posterior polarity by regulating Wnt signaling. Proc. Natl. Acad. Sci. U S A 106, 22329–22334. 10.1073/pnas.0907464106.

28. Scimone, M.L., Cote, L.E., Rogers, T., and Reddien, P.W. (2016). Two FGFRL-Wnt circuits organize the planarian anteroposterior axis. Elife 5. 10.7554/eLife.12845.

29. Lander, R., and Petersen, C.P. (2016). Wnt, Ptk7, and FGFRL expression gradients control trunk positional identity in planarian regeneration. Elife 5. 10.7554/eLife.12850.

30. Cebrià, F., and Newmark, P.A. (2007). Morphogenesis defects are associated with abnormal nervous system regeneration following *roboA* RNAi in planarians. Development 134, 833–837. dev.02794 [pii] 10.1242/dev.02794.

31. Gaviño, M.A., Wenemoser, D., Wang, I.E., and Reddien, P.W. (2013). Tissue absence initiates regeneration through follistatin-mediated inhibition of activin signaling. Elife 2, e00247. 10.7554/eLife.00247.

32. Blassberg, R.A., Felix, D.A., Tejada-Romero, B., and Aboobaker, A.A. (2013). PBX/extradenticle is required to re-establish axial structures and polarity during planarian regeneration. Development 140, 730–739. 10.1242/dev.082982.

33. Tasaki, J., Shibata, N., Nishimura, O., Itomi, K., Tabata, Y., Son, F., Suzuki, N., Araki, R., Abe, M., Agata, K., and Umesono, Y. (2011). ERK signaling controls blastema cell differentiation during planarian regeneration. Development 138, 2417–2427. 10.1242/dev.060764.

34. Sandmann, T., Vogg, M.C., Owlarn, S., Boutros, M., and Bartscherer, K. (2011). The head-regeneration transcriptome of the planarian *Schmidtea mediterranea*. Genome Biol 12, R76. 10.1186/gb-2011-12-8-r76.

35. Wenemoser, D., Lapan, S.W., Wilkinson, A.W., Bell, G.W., and Reddien, P.W. (2012). A molecular wound response program associated with regeneration initiation in planarians. Genes Dev. 26, 988–1002. 10.1101/gad.187377.112.

36. Kao, D., Felix, D., and Aboobaker, A. (2013). The planarian regeneration transcriptome reveals a shared but temporally shifted regulatory program between opposing head and tail scenarios. BMC Genomics 14, 797. 10.1186/1471-2164-14-797.

37. Wurtzel, O., Cote, L.E., Poirier, A., Satija, R., Regev, A., and Reddien, P.W. (2015). A generic and cell-type-specific wound response precedes regeneration in planarians. Dev. Cell 35, 632–645. 10.1016/j.devcel.2015.11.004.

38. Pirotte, N., Stevens, A.S., Fraguas, S., Plusquin, M., Van Roten, A., Van Belleghem, F., Paesen, R., Ameloot, M., Cebrià, F., Artois, T., and Smeets, K. (2015). Reactive Oxygen Species in Planarian Regeneration: An Upstream Necessity for Correct Patterning and Brain Formation. Oxid Med Cell Longev 2015, 392476. 10.1155/2015/392476.

39. Owlarn, S., Klenner, F., Schmidt, D., Rabert, F., Tomasso, A., Reuter, H., Mulaw, M.A., Moritz, S., Gentile, L., Weidinger, G., and Bartscherer, K. (2017). Generic wound signals initiate regeneration in missing-tissue contexts. Nat Commun 8, 2282. 10.1038/s41467-017-02338-x.

40. Jaenen, V., Fraguas, S., Bijnens, K., Heleven, M., Artois, T., Romero, R., Smeets, K., and Cebrià, F. (2021). Reactive oxygen species rescue regeneration after silencing the MAPK-ERK signaling pathway in *Schmidtea mediterranea*. Sci Rep 11, 881. 10.1038/s41598-020-79588-1.

41. González-Sastre, A., De Sousa, N., Adell, T., and Saló, E. (2017). The pioneer factor *Smed-gata456-1* is required for gut cell differentiation and maintenance in planarians. Int. J. Dev. Biol. 61, 53–63. 10.1387/ijdb.160321es.

42. Solari, F., Bateman, A., and Ahringer, J. (1999). The *Caenorhabditis elegans* genes *egl-27* and *egr-1* are similar to MTA1, a member of a chromatin regulatory complex, and are redundantly required for embryonic patterning. Development 126, 2483–2494. 10.1242/dev.126.11.2483.

43. Klein, Y., Halachmi, N., Egoz-Matia, N., Toder, M., and Salzberg, A. (2010). The proprioceptive and contractile systems in *Drosophila* are both patterned by the EGR family transcription factor Stripe. Dev. Biol. 337, 458–470. 10.1016/j.ydbio.2009.11.022.

44. Dussmann, P., Pagel, J.I., Vogel, S., Magnusson, T., Zimmermann, R., Wagner, E., Schaper, W., Ogris, M., and Deindl, E. (2011). Live in vivo imaging of Egr-1 promoter activity during neonatal development, liver regeneration and wound healing. BMC Dev. Biol. 11, 28. 10.1186/1471-213X-11-28.

45. Szwarc, M.M., Hai, L., Gibbons, W.E., Mo, Q., Lanz, R.B., DeMayo, F.J., and Lydon, J.P. (2019). Early growth response 1 transcriptionally primes the human endometrial stromal cell for decidualization. J Steroid Biochem Mol Biol 189, 283–290. 10.1016/j.jsbmb.2019.01.021.

46. Bahrami, S., and Drablos, F. (2016). Gene regulation in the immediate-early response process. Adv Biol Regul 62, 37–49. 10.1016/j.jbior.2016.05.001.

47. Perez-Cadahia, B., Drobic, B., and Davie, J.R. (2011). Activation and function of immediate-early genes in the nervous system. Biochem Cell Biol 89, 61–73. 10.1139/O10-138.

48. Khachigian, L.M. (2006). Early growth response-1 in cardiovascular pathobiology. Circ Res 98, 186–191. 10.1161/01.RES.0000200177.53882.c3.

49. Fan, Y.Y., Ye, G.H., Lin, K.Z., Yu, L.S., Wu, S.Z., Dong, M.W., Han, J.G., Feng, X.P., and Li, X.B. (2013). Time-dependent expression and distribution of Egr-1 during skeletal muscle wound healing in rats. J Mol Histol 44, 75–81. 10.1007/s10735-012-9445-8.

50. Kurosaka, M., Ogura, Y., Funabashi, T., and Akema, T. (2017). Early Growth Response 3 (Egr3) Contributes a Maintenance of C2C12 Myoblast Proliferation. J Cell Physiol 232, 1114–1122. 10.1002/jcp.25574.

51. Magee, N., and Zhang, Y. (2017). Role of early growth response 1 in liver metabolism and liver cancer. Hepatoma Res 3, 268–277. 10.20517/2394-5079.2017.36.

52. Elran, R., Raam, M., Kraus, R., Brekhman, V., Sher, N., Plaschkes, I., Chalifa-Caspi, V., and Lotan, T. (2014). Early and late response of *Nematostella vectensis* transcriptome to heavy metals. Mol. Ecol. 23, 4722–4736. 10.1111/mec.12891.

53. Cary, G.A., Wolff, A., Zueva, O., Pattinato, J., and Hinman, V.F. (2019). Analysis of sea star larval regeneration reveals conserved processes of whole-body regeneration across the metazoa. BMC Biol. 17, 16. 10.1186/s12915-019-0633-9.

54. Gehrke, A.R., Neverett, E., Luo, Y.J., Brandt, A., Ricci, L., Hulett, R.E., Gompers, A., Ruby, J.G., Rokhsar, D.S., Reddien, P.W., and Srivastava, M. (2019). Acoel genome reveals the regulatory landscape of whole-body regeneration. Science 363. 10.1126/science.aau6173.

55. Fraguas, S., Barberán, S., Iglesias, M., Rodríguez-Esteban, G., and Cebrià, F. (2014). *egr-4*, a target of EGFR signaling, is required for the formation of the brain primordia and head regeneration in planarians. Development 141, 1835–1847. 10.1242/dev.101345.

56. Tu, K.C., Cheng, L.C., H, T.K.V., Lange, J.J., McKinney, S.A., Seidel, C.W., and Sánchez Alvarado, A. (2015). Egr-5 is a post-mitotic regulator of planarian epidermal differentiation. Elife 4, e10501. 10.7554/eLife.10501.

57. Robb, S.M., Gotting, K., Ross, E., and Sánchez Alvarado, A. (2015). SmedGD 2.0: The Schmidtea mediterranea genome database. Genesis 53, 535–546. 10.1002/dvg.22872.

58. Grohme, M.A., Schloissnig, S., Rozanski, A., Pippel, M., Young, G.R., Winkler, S., Brandl, H., Henry, I., Dahl, A., Powell, S., et al. (2018). The genome of *Schmidtea mediterranea* and the evolution of core cellular mechanisms. Nature 554, 56–61. 10.1038/nature25473.

59. Rozanski, A., Moon, H., Brandl, H., Martin-Duran, J.M., Grohme, M.A., Huttner, K., Bartscherer, K., Henry, I., and Rink, J.C. (2019). PlanMine 3.0-improvements to a mineable resource of flatworm biology and biodiversity. Nucleic Acids Res. 47, D812–D820. 10.1093/nar/gky1070.

60. Wagner, D.E., Ho, J.J., and Reddien, P.W. (2012). Genetic regulators of a pluripotent adult stem cell system in planarians identified by RNAi and clonal analysis. Cell Stem Cell 10, 299–311. 10.1016/j.stem.2012.01.016.

61. Pearson, B.J., Eisenhoffer, G.T., Gurley, K.A., Rink, J.C., Miller, D.E., and Sánchez Alvarado, A. (2009). Formaldehyde-based whole-mount in situ hybridization method for planarians. Dev. Dyn. 238, 443–450. 10.1002/dvdy.21849.

62. King, R.S., and Newmark, P.A. (2013). *In situ* hybridization protocol for enhanced detection of gene expression in the planarian *Schmidtea mediterranea*. BMC Dev. Biol. 13, 8. 10.1186/1471-213X-13-8.

63. Forsthoefel, D.J., James, N.P., Escobar, D.J., Stary, J.M., Vieira, A.P., Waters, F.A., and Newmark, P.A. (2012). An RNAi screen reveals intestinal regulators of branching morphogenesis, differentiation, and stem cell proliferation in planarians. Dev. Cell 23, 691–704. 10.1016/j.devcel.2012.09.008.

64. Scimone, M.L., Wurtzel, O., Malecek, K., Fincher, C.T., Oderberg, I.M., Kravarik, K.M., and Reddien, P.W. (2018). *foxF-1* Controls Specification of Non-body Wall Muscle and Phagocytic Cells in Planarians. Curr. Biol. 28, 3787–3801 e3786. 10.1016/j.cub.2018.10.030.

65. Benham-Pyle, B.W., Brewster, C.E., Kent, A.M., Mann, F.G., Jr., Chen, S., Scott, A.R., Box, A.C., and Sánchez Alvarado, A. (2021). Identification of rare, transient post-mitotic cell states that are induced by injury and required for whole-body regeneration in *Schmidtea mediterranea*. Nat. Cell Biol. 23, 939–952. 10.1038/s41556-021-00734-6.

66. Scimone, M.L., Cote, L.E., and Reddien, P.W. (2017). Orthogonal muscle fibres have different instructive roles in planarian regeneration. Nature 551, 623–628. 10.1038/nature24660.

67. Scimone, M.L., Cloutier, J.K., Maybrun, C.L., and Reddien, P.W. (2022). The planarian wound epidermis gene *equinox* is required for blastema formation in regeneration. Nat Commun 13, 2726. 10.1038/s41467-022-30412-6.

68. Scimone, M.L., Kravarik, K.M., Lapan, S.W., and Reddien, P.W. (2014). Neoblast specialization in regeneration of the planarian *Schmidtea mediterranea*. Stem Cell Reports 3, 339–352. 10.1016/j.stemcr.2014.06.001.

69. Zeng, A., Li, H., Guo, L., Gao, X., McKinney, S., Wang, Y., Yu, Z., Park, J., Semerad, C., Ross, E., et al. (2018). Prospectively Isolated Tetraspanin(+) Neoblasts Are Adult Pluripotent Stem Cells Underlying Planaria Regeneration. Cell 173, 1593–1608 e1520. 10.1016/j.cell.2018.05.006.

70. Fincher, C.T., Wurtzel, O., de Hoog, T., Kravarik, K.M., and Reddien, P.W. (2018). Cell type transcriptome atlas for the planarian *Schmidtea mediterranea*. Science 360. 10.1126/science.aaq1736.

71. Witchley, J.N., Mayer, M., Wagner, D.E., Owen, J.H., and Reddien, P.W. (2013). Muscle cells provide instructions for planarian regeneration. Cell Rep 4, 633–641. 10.1016/j.celrep.2013.07.022.

72. Vásquez-Doorman, C., and Petersen, C.P. (2014). *zic-1* Expression in planarian neoblasts after injury controls anterior pole regeneration. PLoS Genet. 10, e1004452. 10.1371/journal.pgen.1004452.

73. Currie, K.W., Brown, D.D., Zhu, S., Xu, C., Voisin, V., Bader, G.D., and Pearson, B.J. (2016). HOX gene complement and expression in the planarian *Schmidtea mediterranea*. Evodevo 7, 7. 10.1186/s13227-016-0044-8.

74. Arnold, C.P., Lozano, A.M., Mann, F.G., Jr., Nowotarski, S.H., Haug, J.O., Lange, J.J., Seidel, C.W., and Sánchez Alvarado, A. (2021). Hox genes regulate asexual reproductive behavior and tissue segmentation in adult animals. Nat Commun 12, 6706. 10.1038/s41467-021-26986-2.

75. Newmark, P.A., and Sánchez Alvarado, A. (2000). Bromodeoxyuridine specifically labels the regenerative stem cells of planarians. Dev. Biol. 220, 142–153. 10.1006/dbio.2000.9645.

76. Volohonsky, G., Edenfeld, G., Klambt, C., and Volk, T. (2007). Muscle-dependent maturation of tendon cells is induced by post-transcriptional regulation of *stripeA*. Development 134, 347–356. 10.1242/dev.02735.

77. Hosoda, K., Morimoto, M., Motoishi, M., Nishimura, O., Agata, K., and Umesono, Y. (2016). Simple blood-feeding method for live imaging of gut tube remodeling in regenerating planarians. Dev. Growth Differ. 58, 260–269. 10.1111/dgd.12270.

78. Kellermeyer, R., Heydman, L.M., Gillis, T., Mastick, G.S., Song, M., and Kidd, T. (2020). Proteolytic cleavage of Slit by the Tolkin protease converts an axon repulsion cue to an axon growth cue in vivo. Development 147. 10.1242/dev.196055.

79. DuBuc, T.Q., Traylor-Knowles, N., and Martindale, M.Q. (2014). Initiating a regenerative response; cellular and molecular features of wound healing in the cnidarian *Nematostella vectensis*. BMC Biol. 12, 24. 10.1186/1741-7007-12-24.

80. De Simone, A., Evanitsky, M.N., Hayden, L., Cox, B.D., Wang, J., Tornini, V.A., Ou, J., Chao, A., Poss, K.D., and Di Talia, S. (2021). Control of osteoblast regeneration by a train of Erk activity waves. Nature 590, 129–133. 10.1038/s41586-020-03085-8.

81. Fraguas, S., Umesono, Y., Agata, K., and Cebria, F. (2017). Analyzing pERK Activation During Planarian Regeneration. Methods Mol Biol 1487, 303–315. 10.1007/978-1-4939-6424-6_23.

82. Bohr, T.E., Shiroor, D.A., and Adler, C.E. (2021). Planarian stem cells sense the identity of the missing pharynx to launch its targeted regeneration. Elife 10. 10.7554/eLife.68830.

83. Santiago, F.S., Lowe, H.C., Kavurma, M.M., Chesterman, C.N., Baker, A., Atkins, D.G., and Khachigian, L.M. (1999). New DNA enzyme targeting Egr-1 mRNA inhibits vascular smooth muscle proliferation and regrowth after injury. Nat. Med. 5, 1264–1269. 10.1038/15215.

84. Dieckgraefe, B.K., and Weems, D.M. (1999). Epithelial injury induces Egr-1 and Fos expression by a pathway involving protein kinase C and ERK. Am J Physiol 276, G322–330. 10.1152/ajpgi.1999.276.2.G322.

85. Dhara, S.P., Rau, A., Flister, M.J., Recka, N.M., Laiosa, M.D., Auer, P.L., and Udvadia, A.J. (2019). Cellular reprogramming for successful CNS axon regeneration is driven by a temporally changing cast of transcription factors. Sci Rep 9, 14198. 10.1038/s41598-019-50485-6.

86. Kang, J., Hu, J., Karra, R., Dickson, A.L., Tornini, V.A., Nachtrab, G., Gemberling, M., Goldman, J.A., Black, B.L., and Poss, K.D. (2016). Modulation of tissue repair by regeneration enhancer elements. Nature 532, 201–206. 10.1038/nature17644.

87. Qian, L., Huang, Y., Spencer, C.I., Foley, A., Vedantham, V., Liu, L., Conway, S.J., Fu, J.D., and Srivastava, D. (2012). *In vivo* reprogramming of murine cardiac fibroblasts into induced cardiomyocytes. Nature 485, 593–598. 10.1038/nature11044.

88. Kurita, M., Araoka, T., Hishida, T., O’Keefe, D.D., Takahashi, Y., Sakamoto, A., Sakurai, M., Suzuki, K., Wu, J., Yamamoto, M., et al. (2018). In vivo reprogramming of wound-resident cells generates skin epithelial tissue. Nature 561, 243–247. 10.1038/s41586-018-0477-4.

89. Guo, Z., Zhang, L., Wu, Z., Chen, Y., Wang, F., and Chen, G. (2014). In vivo direct reprogramming of reactive glial cells into functional neurons after brain injury and in an Alzheimer’s disease model. Cell Stem Cell 14, 188–202. 10.1016/j.stem.2013.12.001.

90. Grobstein, C. (1956). Inductive tissue interaction in development. Adv. Cancer Res. 4, 187–236. 10.1016/s0065-230x(08)60725-3.

91. Dressler, G.R. (2006). The cellular basis of kidney development. Annu. Rev. Cell. Dev. Biol. 22, 509–529. 10.1146/annurev.cellbio.22.010305.104340.

92. Sai, X., and Ladher, R.K. (2015). Early steps in inner ear development: induction and morphogenesis of the otic placode. Front Pharmacol 6, 19. 10.3389/fphar.2015.00019.

93. Henry, J.J., and Tsonis, P.A. (2010). Molecular and cellular aspects of amphibian lens regeneration. Prog Retin Eye Res 29, 543–555. 10.1016/j.preteyeres.2010.07.002.

94. Talman, V., and Kivelä, R. (2018). Cardiomyocyte-Endothelial Cell Interactions in Cardiac Remodeling and Regeneration. Front Cardiovasc Med 5, 101. 10.3389/fcvm.2018.00101.

95. Bassat, E., and Tanaka, E.M. (2021). The cellular and signaling dynamics of salamander limb regeneration. Curr. Opin. Cell Biol. 73, 117–123. 10.1016/j.ceb.2021.07.010.

96. Koike, H., Manabe, I., and Oishi, Y. (2022). Mechanisms of cooperative cell-cell interactions in skeletal muscle regeneration. Inflamm Regen 42, 48. 10.1186/s41232-022-00234-6.

97. Lancaster, M.A., and Knoblich, J.A. (2014). Organogenesis in a dish: modeling development and disease using organoid technologies. Science 345, 1247125. 10.1126/science.1247125.

98. Hofer, M., and Lutolf, M.P. (2021). Engineering organoids. Nat Rev Mater 6, 402–420. 10.1038/s41578-021-00279-y.

99. Lubarsky, B., and Krasnow, M.A. (2003). Tube morphogenesis: making and shaping biological tubes. Cell 112, 19–28. 10.1016/s0092-8674(02)01283-7.

100. Iruela-Arispe, M.L., and Beitel, G.J. (2013). Tubulogenesis. Development 140, 2851–2855. 10.1242/dev.070680.

101. Londono, R., and Badylak, S.F. (2015). Regenerative Medicine Strategies for Esophageal Repair. Tissue Eng Part B Rev 21, 393–410. 10.1089/ten.TEB.2015.0014.

102. Sánchez Alvarado, A., Newmark, P.A., Robb, S.M., and Juste, R. (2002). The *Schmidtea mediterranea* database as a molecular resource for studying platyhelminthes, stem cells and regeneration. Development 129, 5659–5665. 10.1242/dev.00167.

103. Roberts-Galbraith, R.H., and Newmark, P.A. (2013). Follistatin antagonizes Activin signaling and acts with Notum to direct planarian head regeneration. Proc Natl Acad Sci U S A 110, 1363–1368. 10.1073/pnas.1214053110.

104. Vila-Farré, M., and Rink, J.C. (2018). The Ecology of Freshwater Planarians. Methods Mol Biol 1774, 173–205. 10.1007/978-1-4939-7802-1_3.

105. Reddien, P.W., Newmark, P.A., and Sanchez Alvarado, A. (2008). Gene nomenclature guidelines for the planarian Schmidtea mediterranea. Dev Dyn 237, 3099–3101. 10.1002/dvdy.21623.

106. Guindon, S., Dufayard, J.F., Lefort, V., Anisimova, M., Hordijk, W., and Gascuel, O. (2010). New algorithms and methods to estimate maximum-likelihood phylogenies: assessing the performance of PhyML 3.0. Syst. Biol. 59, 307–321. 10.1093/sysbio/syq010.

107. Lefort, V., Longueville, J.E., and Gascuel, O. (2017). SMS: Smart Model Selection in PhyML. Mol. Biol. Evol. 34, 2422–2424. 10.1093/molbev/msx149.

108. Zayas, R.M., Hernandez, A., Habermann, B., Wang, Y., Stary, J.M., and Newmark, P.A. (2005). The planarian *Schmidtea mediterranea* as a model for epigenetic germ cell specification: analysis of ESTs from the hermaphroditic strain. Proc. Natl. Acad. Sci. USA 102, 18491–18496.

109. Collins, J.J., 3rd, Hou, X., Romanova, E.V., Lambrus, B.G., Miller, C.M., Saberi, A., Sweedler, J.V., and Newmark, P.A. (2010). Genome-wide analyses reveal a role for peptide hormones in planarian germline development. PLoS Biol. 8, e1000509. 10.1371/journal.pbio.1000509.

110. Forsthoefel, D.J., Cejda, N.I., Khan, U.W., and Newmark, P.A. (2020). Cell-type diversity and regionalized gene expression in the planarian intestine. Elife 9. 10.7554/eLife.52613.

111. Rouhana, L., Weiss, J.A., Forsthoefel, D.J., Lee, H., King, R.S., Inoue, T., Shibata, N., Agata, K., and Newmark, P.A. (2013). RNA interference by feeding in vitro-synthesized double-stranded RNA to planarians: methodology and dynamics. Dev. Dyn. 242, 718–730. 10.1002/dvdy.23950.

112. Ross, K.G., Omuro, K.C., Taylor, M.R., Munday, R.K., Hubert, A., King, R.S., and Zayas, R.M. (2015). Novel monoclonal antibodies to study tissue regeneration in planarians. BMC Dev. Biol. 15, 2. 10.1186/s12861-014-0050-9.

113. Brandl, H., Moon, H., Vila-Farre, M., Liu, S.Y., Henry, I., and Rink, J.C. (2016). PlanMine--a mineable resource of planarian biology and biodiversity. Nucleic Acids Res. 44, D764–773. 10.1093/nar/gkv1148.

114. Andrews, S. (2010). FastQC: a quality control tool for high throughput sequence data.

115. Langmead, B., and Salzberg, S.L. (2012). Fast gapped-read alignment with Bowtie 2. Nat. Methods 9, 357–359. 10.1038/nmeth.1923.

116. Liao, Y., Smyth, G.K., and Shi, W. (2019). The R package Rsubread is easier, faster, cheaper and better for alignment and quantification of RNA sequencing reads. Nucleic Acids Res. 47, e47. 10.1093/nar/gkz114.

117. R Core Team (2021). R: A language and environment for statistical computing. URL https://www.R-project.org/.

118. Robinson, M.D., McCarthy, D.J., and Smyth, G.K. (2010). edgeR: a Bioconductor package for differential expression analysis of digital gene expression data. Bioinformatics 26, 139–140. 10.1093/bioinformatics/btp616.

119. Hao, Y., Hao, S., Andersen-Nissen, E., Mauck, W.M., 3rd, Zheng, S., Butler, A., Lee, M.J., Wilk, A.J., Darby, C., Zager, M., et al. (2021). Integrated analysis of multimodal single-cell data. Cell 184, 3573–3587 e3529. 10.1016/j.cell.2021.04.048.

120. Hafemeister, C., and Satija, R. (2019). Normalization and variance stabilization of single-cell RNA-seq data using regularized negative binomial regression. Genome Biol 20, 296. 10.1186/s13059-019-1874-1.

121. Stuart, T., Butler, A., Hoffman, P., Hafemeister, C., Papalexi, E., Mauck, W.M., 3rd, Hao, Y., Stoeckius, M., Smibert, P., and Satija, R. (2019). Comprehensive Integration of Single-Cell Data. Cell 177, 1888–1902 e1821. 10.1016/j.cell.2019.05.031.

122. Plass, M., Solana, J., Wolf, F.A., Ayoub, S., Misios, A., Glazar, P., Obermayer, B., Theis, F.J., Kocks, C., and Rajewsky, N. (2018). Cell type atlas and lineage tree of a whole complex animal by single-cell transcriptomics. Science 360. 10.1126/science.aaq1723.

123. Livak, K.J., and Schmittgen, T.D. (2001). Analysis of relative gene expression data using real-time quantitative PCR and the 2(-Delta Delta C(T)) Method. Methods 25, 402–408. 10.1006/meth.2001.1262.

124. Schneider, C.A., Rasband, W.S., and Eliceiri, K.W. (2012). NIH Image to ImageJ: 25 years of image analysis. Nat. Methods 9, 671–675. 10.1038/nmeth.2089

